# High-throughput screen identifies a potent MCU activator boosting cardiac contractile bioenergetics

**DOI:** 10.64898/2026.09.04.749429

**Authors:** Elena Caldero-Escudero, Marilén Federico, Angel Sanchez-Gonzalez, Adria Gil, Silvia Fernandez-Martinez, Paloma García-Casas, Rosalba I Fonteriz, Flavien Bermont, Benjamin Brinon, Silvia Romero-Sanz, Andrea Moran-Cerro, Luis Gonzalez-Moreno, Araceli del Arco, Shey-Shing Sheu, Jerome N Feige, Mayte Montero, Javier Alvarez, Umberto de Marchi, Sergio De la Fuente, Jaime Santo-Domingo

## Abstract

Mitochondrial Ca^2+^ uptake is an evolutionarily conserved mechanism mediated by the mitochondrial calcium uniporter complex (MCUc) that couples intracellular Ca²⁺ signaling to energy metabolism and cellular function. Despite its physiological importance, potent and selective pharmacological activators of the MCU complex remain scarce, and existing compounds show modest specificity or poorly explored mechanisms. Here, we performed a high-content screen of 1,280 bioactive compounds and identified CGP7930 as a potent activator of mitochondrial Ca²⁺ uptake. Mechanistically, the compound required MICU1, but not MICU2, to exert its effects. Molecular docking, analysis of the topology of electron density (ρ) to explore the Non-Covalent Interaction (NCI) index, and site-directed mutagenesis identified a critical interaction site within MICU1 involving residues Gln304 and Val307. In addition to directly stimulating MCU activity, CGP7930 increased mitochondria-endoplasmic reticulum contact sites, suggesting an additional mechanism to facilitate inter-organellar Ca²⁺ transfer. CGP7930 promoted Ca^2+^-dependent activation of mitochondrial energy metabolism in cardiomyocytes and boosted the contractile performance of the mice hearts. This effect was MCU-dependent, as hearts from MCU KO mice failed to increase the ventricular contraction force upon CGP7930 perfusion. We conclude that MICU1 is a druggable regulatory target for modulating mitochondrial Ca²⁺ signaling, and that CGP7930 is a new potent activator of the MCU that promotes excitation-bioenergetics-contraction coupling in the heart.

**HIGHLIGTS:**

- We screen a library containing 1280 compounds with well-defined targets to identify new potential mechanisms activating ER to mitochondria Ca^2+^ transport.
- CGP7930 is a new direct and potent activator of the mitochondrial Ca^2+^ uniporter.
- CGP7930 binds MICU1 on a specific binding pocket located at the cleft between the N and C lobes and coordinated by Glu-304 and Val-307.
- Pharmacological activation of MCU with CGP7930 remodels ER Ca^2+^ homeostasis and ER-mitochondria contact sites.
- Pharmacological activation of MCU with CGP7930 boosts excitation-bioenergetics-contraction coupling in the heart.

## INTRODUCTION

Calcium ion (Ca²⁺) is a ubiquitous second messenger that regulates a wide range of cellular processes, including metabolism, gene expression, secretion, proliferation, and cell death^1^. Among intracellular organelles, mitochondria play a central role in decoding cytosolic Ca²⁺ signals and translating them into metabolic responses. Transient increases in mitochondrial matrix Ca²⁺ stimulate key dehydrogenases of the tricarboxylic acid (TCA) cycle, thereby enhancing oxidative phosphorylation and ATP production to match cellular energy demand^2–4^. Beyond its role in bioenergetics, mitochondrial Ca²⁺ signaling contributes to the regulation of reactive oxygen species (ROS) generation, intracellular signaling pathways, and cell fate decisions^5^. Conversely, excessive mitochondrial Ca²⁺ accumulation can trigger permeability transition pore opening, mitochondrial dysfunction, and cell death, implicating mitochondrial Ca²⁺ homeostasis in numerous pathological conditions, including cardiovascular diseases, neurodegeneration, cancer, and metabolic disorders^6,7^.

Mitochondrial Ca²⁺ uptake is mediated by the mitochondrial calcium uniporter (MCU) complex, a highly selective channel located in the inner mitochondrial membrane^8–10^. The pore-forming subunit MCU assembles into a tetrameric structure that conducts Ca²⁺ into the mitochondrial matrix, driven by the large electrochemical gradient across the inner membrane^11^. MCU activity is tightly regulated by several auxiliary proteins, including the essential MCU regulator^12^ (EMRE) and the EF-hand-containing proteins MICU1, MICU2, and MICU3^13^. Under resting cytosolic Ca²⁺ concentrations, MICU proteins maintain the channel in a closed or low-conductance state, preventing excessive mitochondrial Ca²⁺ accumulation. Upon cytosolic Ca²⁺ elevation, Ca²⁺ binding to MICU proteins relieves this inhibition, promoting channel opening and cooperative activation of mitochondrial Ca²⁺ uptake^14^. This sophisticated gating mechanism allows mitochondria to respond selectively to physiological Ca²⁺ signals while avoiding Ca²⁺ overload^15^.

The relatively low affinity of the MCU complex for Ca²⁺ requires mitochondria to be exposed to local high Ca²⁺ microdomains in order to achieve efficient uptake^16^. These microdomains are generated at specialized contact sites between the endoplasmic reticulum (ER) and mitochondria, commonly referred to as mitochondria-ER contact sites^17^ (MERCS). At these interfaces, Ca²⁺ released through inositol 1,4,5-trisphosphate receptors (IP₃Rs) or ryanodine receptors reaches concentrations substantially higher than those found in the bulk cytosol, thereby efficiently activating the MCU complex^18^. Beyond facilitating Ca²⁺ transfer, MERCS coordinate multiple cellular functions, including lipid exchange, mitochondrial dynamics, autophagy, apoptosis, and metabolic adaptation_19_. Alterations in MERCS architecture or function have been associated with numerous human diseases, highlighting the importance of ER-mitochondria communication in cellular physiology and pathophysiology^20,21^. For instance, this spatial arrangement is essential during excitation-contraction coupling in muscle tissue, as it allows mitochondria to override the MCU’s gatekeeping threshold instantly, coupling the metabolic needs of contraction directly to rapid ATP production via oxidative metabolism^22^.

Given the central role of mitochondrial Ca²⁺ signaling in cell physiology and disease, considerable efforts have been devoted to identifying pharmacological modulators of the MCU complex. Early inhibitors such as Ruthenium Red and Ru360 provided invaluable experimental tools but were limited by poor cell permeability and unfavorable pharmacological properties. More recently, a permeable Ru265 was developed, which binds to and blocks the selectivity filter at the mouth of the channel^23^. Other negative modulators include the antineoplastic drug mitoxantrone, which targets the acidic residues of the MCU selectivity filter^24^, and the MICU1-binding compounds MCU-i4 and MCU-i11, which decrease mitochondrial Ca^2+^ influx by stabilizing the gatekeeping state of the complex^25^. Inhibitors have demonstrated therapeutic potential in preclinical settings ranging from ischemia-reperfusion injury and neurodegeneration to cancer^26–28^. Conversely, MCU activators are pursued to treat conditions involving bioenergetic failure. Polyamines such as spermine and plant flavonoids such as kaempferol^29^ have been shown to sensitize the MCU to Ca^2+^. More recently, the morpholine derivative amorolfine^30^ and the olive-derived polyphenol oleuropein have been identified as potent positive modulators. Oleuropein, in particular, binds specifically to a cleft in the MICU1 subunit and has been shown to stimulate mitochondrial respiration and ATP synthesis by enhancing Ca^2+^ entry, effectively reversing age-related bioenergetic decline in skeletal muscle cells across species^31^.

Nevertheless, the number of selective MCU activators remains limited, and the molecular mechanisms underlying their activity are often incompletely understood. In the present study, using a high-content screening workflow, we identify and characterize CGP7930 as a novel potent activator of the MCU complex, define its mechanism of action, and demonstrate its ability to enhance mitochondrial Ca²⁺ uptake and ER–mitochondria communication. We further demonstrate that CGP7930 improved cardiac contractile function by enhancing energetic coupling in an MCU-dependent manner.

## RESULTS

### High-content screening workflow identifies CGP7930 as a new potent pharmacological modulator of mitochondrial Ca^2+^ uptake

To identify novel regulatory mechanisms involved in ER-to-mitochondria Ca²⁺ transfer (Fig. 1A), we screened the TOCRISCREEN® library (Fig. 1B), comprising 1,280 biologically active compounds with well-defined molecular targets, for their ability to modulate histamine-induced mitochondrial Ca²⁺ uptake in HeLa cells expressing a mitochondria-targeted aequorin (mitmutAEQ) and reconstituted with coelenterazine (Fig. 1A-B). The library was screened across four assay plates (A–D), using kaempferol, a previously-described modulator of mitochondrial Ca²⁺ uptake, as a positive control (Fig. 1C). For each compound, the response was normalized to the average signal obtained in vehicle-treated control cells (set to 0%; blue) and to the average response elicited by kaempferol (set to 100%; red). A threshold of 50% activation relative to kaempferol was established (Fig. 1C, cyan line), and compounds exceeding this value were classified as potential positive modulators of ER-to-mitochondria Ca²⁺ transfer (Fig. 1C, cyan outline with green fill). Using these criteria, we identified 18 primary hits, corresponding to a hit rate of 1.41% (Fig. 1C). Compounds below this threshold were considered negative (Fig. 1C, green). To validate these findings, the 18 primary hits were subjected to a reconfirmation screen, which verified 11 compounds exhibiting activation levels above 50%, including six compounds that exceeded the response induced by kaempferol (>100% activation) (Fig. 1D). The confirmed hits were subsequently evaluated in a counter-screen designed to determine whether they selectively enhanced mitochondrial Ca²⁺ uptake without inducing a concomitant increase in cytosolic Ca²⁺ levels (Fig. 1B and E). This analysis excluded three compounds, resulting in a final set of five positive hits that displayed greater activity than kaempferol without affecting cytosolic Ca^2+^ (Supplementary Table 1).

**Fig. 1.**
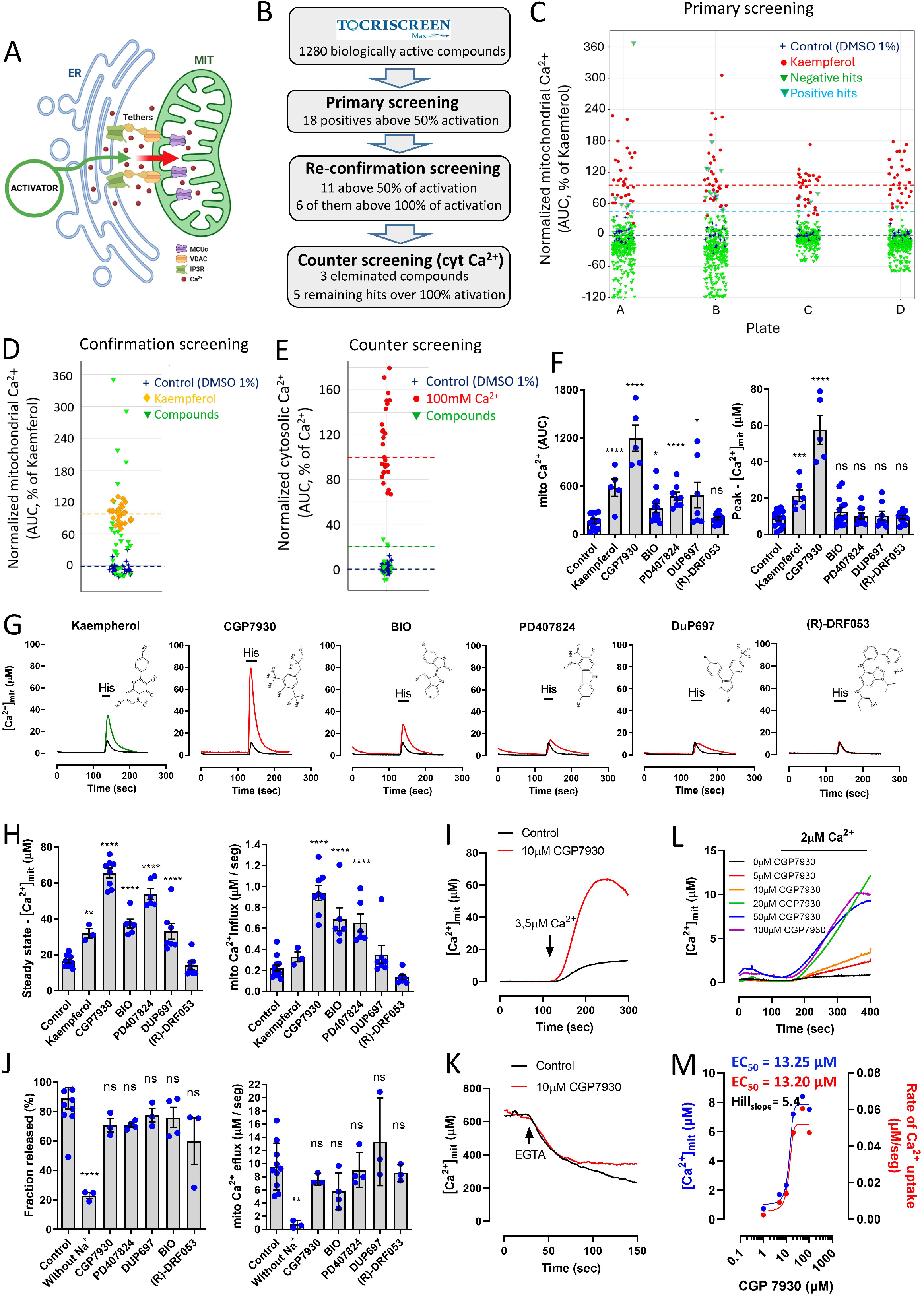
High content screening and validation of candidate compounds that enhance mitochondrial Ca²⁺ uptake in intact and permeabilized cells. (A) Schematic representation. The screening aims to identify compounds and potential targets that enhance ER to mitochondrial Ca2+ transport upon IP3R activation by histamine in HeLa cells. (B) Workflow and results of the screening. (C) Results of mtCa2+ primary screen in HeLa cells stimulated with 100 µM histamine; The TOCRISCREEN Max® library was measured in 4 plates (A/B/C/D). The effect of the hits was normalized in the figure on the average of control cells (set to 0, blue) and on the average of positive control cells set to 100 (Kaempferol, red). The established threshold is represented in cyan; all the hits below the selected threshold are considered negative (green); all the hits above the threshold are considered positive (cyan border, green fill). (D) Reconfirmation of positive hits from the primary screen using the same assay. Positive control, kaempferol 10 µM. (E) Counter-screen of cytosolic Ca2+ to exclude the hits that modulate both mtCa2+ and cytosolic Ca2+; positive control is 100 mM extracellular Ca2+. (F) Validation of the 5 primary hits of the screening in a small-scale perfusion system. Impact of hits on mitochondrial Ca²⁺ uptake in response to histamine stimulation, expressed as area under the curve (AUC) and peak mitochondrial Ca²⁺ concentration ([Ca²⁺]ₘ peak). Data are presented as mean ± s.e.m. from 5-15 independent experiments per condition. (G) Representative traces of [Ca²⁺]ₘ evoked by stimulation with 100 μM histamine in HeLa cells preincubated with kaempferol, PD407824, DuP697, BIO, CGP7930, and (R)-DRF053. (H) Quantification of steady-state [Ca²⁺]ₘ and the quantification of the initial mitochondrial Ca²⁺ uptake rate under the indicated conditions. Data are presented as mean ± s.e.m. from 3-10 independent experiments per condition. (I) Representative time courses of [Ca²⁺]ₘ. Mitochondrial Ca²⁺ uptake in permeabilized HeLa cells was induced by perfusion with intracellular-like medium containing 3.5 μM free Ca²⁺. (J) Quantification of the total fraction of mitochondrial Ca²⁺ released under the indicated conditions, expressed as a percentage of steady-state [Ca²⁺]ₘ and initial mitochondrial Ca²⁺ efflux rate. Data are presented as mean ± s.e.m. from 3-8 independent experiments per condition. (K) Representative time courses of [Ca²⁺]ₘ measured in permeabilized HeLa cells. After mitochondrial Ca²⁺ uptake reached a steady-state level, 10 μM EGTA was added to the perfusion buffer to induce Ca²⁺ release. (L) Representative time courses of [Ca²⁺]ₘ measured in permeabilized HeLa cells. Mitochondrial Ca²⁺ uptake was induced by perfusion with 2 μM free Ca²⁺ and increasing concentrations of CGP7930, from 0 to 100 μM were tested. (M) Dose-response effect of CGP7930 in permeabilized cells perfused with intracellular-like medium containing 2 μM free Ca²⁺.

To further validate these findings, we verified the impact of the 5 remaining hits with newly synthesized compounds on histamine-induced mitochondrial Ca^2+^ uptake. We used a small-scale perfusion system that enables analysis of rapid changes in Ca^2+^ dynamics following plasma-membrane receptor activation using the bioluminescent Ca²⁺ sensor mitmutAEQ (Fig. 1G). To quantify the effects of each compound on histamine-induced mitochondrial Ca²⁺ dynamics, we estimated the total Ca^2+^ mobilized, calculated as the area under the curve (AUC) of the resulting Ca²⁺ transients (Fig. 1G). As expected, kaempferol increased the AUC by approximately threefold compared to control conditions (Fig. 1F-G). Similarly, CGP7930, BIO, PD407824, and DuP697 all produced larger Ca²⁺ transients than control, with CGP7930 showing the strongest effect, yielding an approximately fivefold increase in AUC (Fig. 1F). In contrast, (R)-DRF053, although identified in the initial screen, did not significantly alter the AUC relative to control (Fig. 1F-G). We next analyzed the height of mitochondrial Ca^2+^ concentration ([Ca²⁺]ₘ) peak elicited by histamine. Among the 5 screening-filtered compounds, only CGP7930 significantly increased [Ca²⁺]ₘ peak values compared to the control condition. Kaempferol induced an approximately threefold increase, whereas CGP7930 elicited a ∼sixfold enhancement. (Fig. 1F-G).

Collectively, the reported molecular targets of these compounds do not suggest an obvious connection with intracellular Ca²⁺ homeostasis (Supplementary Table 1). Regardless of its previously defined targets, compound-driven elevations in [Ca²⁺]ₘ may arise from enhanced ER Ca²⁺ release, increased ER-mitochondrial physical coupling, or direct modulation of mitochondrial Ca²⁺ transport systems. To investigate the latter possibility, mitochondrial Ca²⁺ uptake was measured in permeabilized HeLa cells, a model that allows selective assessment of mitochondrial Ca²⁺ influx. Permeabilized cells were exposed to a buffered solution containing 3.5 μM Ca²⁺ in the presence of each compound, and both steady-state [Ca²⁺]ₘ and uptake rates were determined (Fig. 1H and Supplementary Fig.1A). All compounds, except (R)-DRF053, significantly increased steady-state [Ca²⁺]ₘ compared to control, consistent with enhanced MCUc activity. Analysis of mitochondrial Ca²⁺ influx rates revealed again that all tested compounds, except for (R)-DRF053, accelerated Ca²⁺ uptake. Among the tested compounds, CGP7930 consistently exhibited the strongest effect, producing both the highest steady-state [Ca²⁺]ₘ and the fastest Ca²⁺ influx rate (Fig. 1H-I and Supplementary Fig.1A). Given that uptake rates and steady-state [Ca²⁺]ₘ reflect the balance between Ca²⁺ entry through the MCUc and extrusion via Na⁺ and H^+^ exchangers, we next examined mitochondrial Ca²⁺ release dynamics. Permeabilized HeLa cells were first loaded with Ca²⁺ by controlled perfusion of 5 µM Ca²⁺, and once the steady state was reached, the cells were perfused with the Ca²⁺ chelator EGTA to isolate Ca²⁺ release fluxes. Removal of extramitochondrial Na⁺ effectively abolished Ca²⁺ extrusion, confirming that Ca²⁺ release under these conditions is predominantly mediated by NCLX (Fig. 1J and Supplementary Fig.1B). In contrast, none of the tested compounds altered either the initial rate of mitochondrial Ca²⁺ efflux or the percentage of Ca²⁺ released compared to control conditions (Fig. 1J and Supplementary Fig.1B). These results indicate that the compounds identified in our screen do not measurably affect mitochondrial Ca²⁺ efflux, supporting the conclusion that their primary mode of action is to enhance Ca²⁺ influx.

HeLa cells permeabilized and perfused with an intracellular Ca^2+^ buffer containing 2 μM Ca^2+^ (Fig. 1L) showed that CGP7930 dose-dependently activated [Ca²⁺]ₘ uptake. EC_50_ values were 13.20 µM when steady state values were considered, or 13.25 µM for Ca^2+^ uptake rates (Fig.1L-M). Non-linear regression analysis of the dose-response data yielded a remarkably steep Hill coefficient (nH ̴̴ 5), indicating a high degree of positive cooperativity. This value aligns with the known macromolecular architecture of the MCUc and its key regulatory subunits, MICU1 and MICU2. Taken together, our data demonstrate that the validated compounds selectively promote mitochondrial Ca²⁺ uptake by directly or indirectly stimulating MCUc activity, without altering NCLX-dependent Ca²⁺ extrusion. Among them, CGP7930 consistently exhibited the most pronounced effects across all experimental conditions, producing the largest increases in Ca²⁺ uptake, steady-state [Ca²⁺]ₘ, and mitochondrial Ca²⁺ influx rates. Based on its robust and reproducible enhancement of MCUc-dependent Ca²⁺ entry, CGP7930 was selected as the lead compound for subsequent mechanistic and functional studies.

### Pharmacological activation of MCUc with CGP7930 requires the regulatory subunit MICU1

The mitochondrial calcium uniporter is a high-molecular-weight complex (MCUc) composed of (Fig.2A): MCU, the pore-forming tetrameric channel in the inner mitochondrial membrane that conducts Ca²⁺ into the matrix; MCUb, a dominant-negative paralog that lowers conductance; MICU1 and MICU2, EF-hand Ca²⁺-sensing regulatory subunits on the intermembrane space side that set the activation threshold and gate MCU activity; EMRE, a small single-pass membrane protein required for structural coupling of MICUs to MCU and for efficient uniporter function; and MCUR, which binds to MCU and EMRE and functions as a scaffold that regulates MCUc activity. Here, we have systematically explored the subunit requirements for MCUc CGP7930 activation. Accordingly, we KO each component of the channel in HAP1 cells and measured steady-state [Ca²⁺]ₘ and uptake rates in permeabilized cells perfused with low micromolar intracellular Ca^2+^ buffers in the absence or presence of CGP7930.

**Fig. 2.**
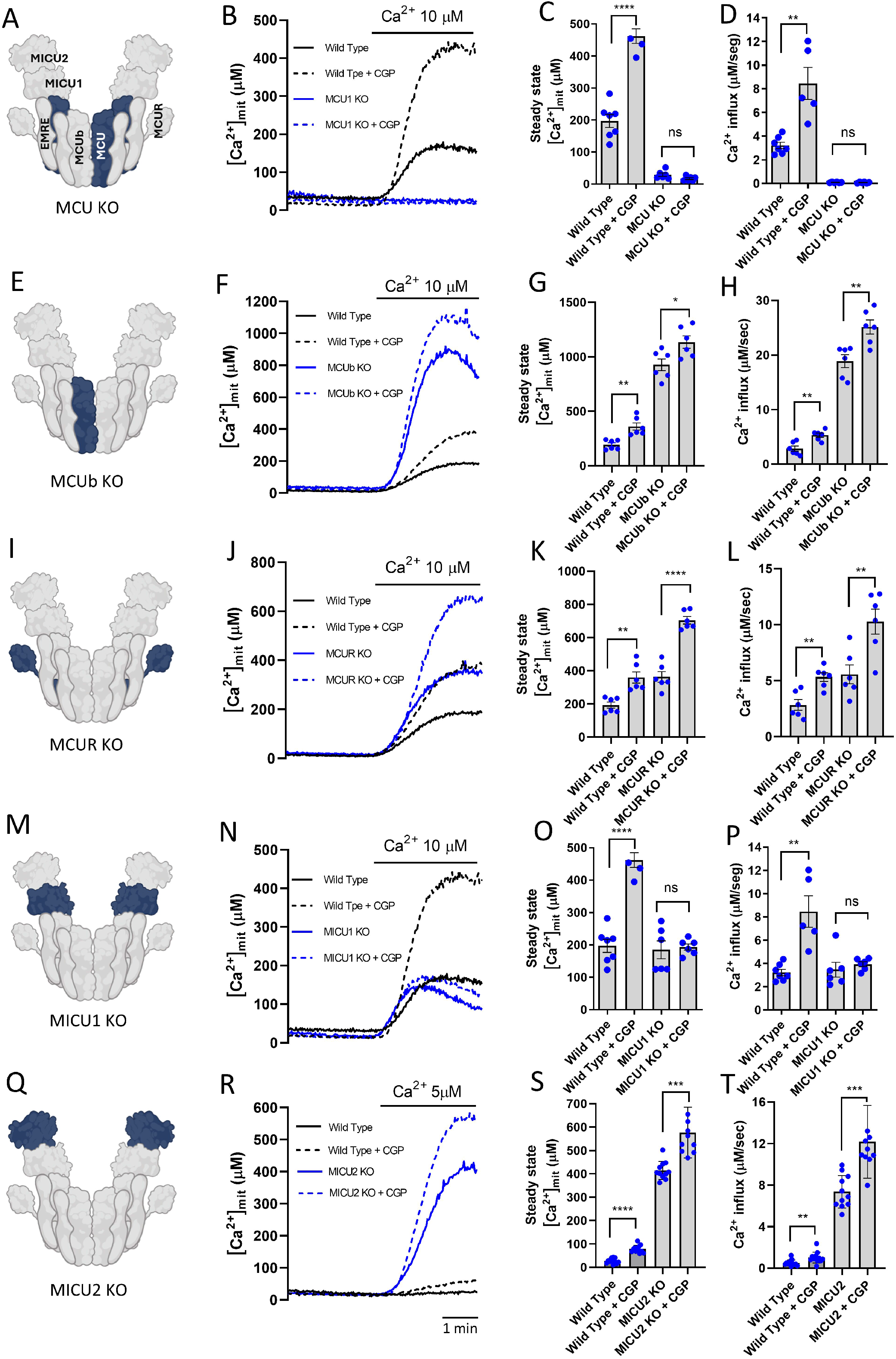
Effect of systematic loss of function of MCUc subunits on CGP7930-induced mitochondrial Ca2+ uptake. (A, E, I, M, Q) Schematic representation of the MCUc, including all the subunits tested. KO subunit in each experiment is displayed in blue. (B, F, J, N, R) Effect of 10 CGP7930 on the average time courses of [Ca²⁺]ₘ measured in permeabilized wild-type HAP1 cells and MCUc-subunits KO HAP1 cells expressing mitochondria-targeted single-mutant aequorin (mit-119mutAEQ) and reconstituted with coelenterazine w. Mitochondrial Ca²⁺ uptake was induced by perfusion with intracellular-like medium containing 10 or 5 μM free Ca²⁺. (C, G, K, O, S) Quantification of steady state [Ca²⁺]ₘ reached under each experimental condition. (D, H, L, P, T) Quantification of the initial mitochondrial Ca²⁺ uptake rate under the indicated conditions. Data are presented as mean ± s.e.m. from 3-11 independent experiments per condition.

Consistent with the stimulatory effect on Ca^2+^ uptake in HeLa cells, 10µM CGP7930 also increased mitochondrial Ca^2+^ uptake in HAP1 wild-type cells (Fig.2B-D). As previously reported, the MCU loss of function abolished mitochondrial Ca^2+^ uptake (Fig.2B-D). CGP7930 did not rescue Ca^2+^ transport in MCU loss-of-function cells, suggesting that CGP7930 requires the main pore-forming subunit of MCUc, and thus does not engage any of the MCU-independent mechanisms reported for mitochondrial Ca^2+^ uptake^32^. We further explored the possibility that CGP7930 action relies on the negative-modulator pore-forming subunit MCUb (Fig.2E). As previously reported, MCUb loss of function also increased the ability of MCUc to permeate Ca^2+^ (Fig.2F-H). However, lack of MCUb did not alter the stimulatory effect of CGP7930, suggesting that this subunit is not required for CGP7930-induced activation of MCUc. We then tested whether the activation properties of CGP7930 were mediated by the scaffold subunit MCUR1^33^ (Fig.2I). Contrary to previous reports, in our hands MCUR1 loss of function also increased the ability of MCUc to permeate Ca^2+^ (Fig.1J-L). However, lack of MCUR1 did not alter the stimulatory effect of CGP7930 (Fig.1J-L), suggesting that this subunit is not required for CGP7930-induced activation of MCUc.

Subsequently, we explored whether the MICU1 and MICU2 regulatory subunits mediated the effect of CGP7930 on [Ca²⁺]ₘ uptake (Fig.2M and Q). As previously reported^14,34^, MICU1 loss of function in HAP1 cells permeabilized and perfused with a 10 µM Ca^2+^ buffer did not alter the kinetics of Ca^2+^ uptake (Fig. 2M-P). However, the lack of MICU1 abolished the stimulatory effect of CGP7930, suggesting that this subunit is required for CGP7930-induced activation of MCUc. Interestingly, the lack of MICU1 is associated with concomitant loss of the MICU2 subunit^35^. Therefore, to distinguish the contributions of both subunits, we also tested the impact of CGP7930 on MICU2 KO HAP1 cells (Fig. 2R-T). As previously reported in HeLa cells^36^, MICU2 loss of function in HAP1 cells also increased Ca^2+^ uptake at low micromolar Ca^2+^ levels. However, lack of MICU2 did not impact the stimulatory effect of CGP7930, suggesting that this subunit is not required for CGP7930-induced activation of MCUc. Collectively, the results of this functional genetic analysis indicate that the activation property of CGP7930 on the capacity of the MCUc to transport Ca^2+^ is mediated by MICU1.

### Pharmacological activation of MCUc with CGP7930 relies on the Gln-304 and Val-307 located at the cleft between the C- and N-lobes of MICU1

Having identified MICU1 as the MCUc subunit mediating the effects of CGP7930, we next sought to define its binding site in greater detail. To this end, we performed computational docking simulations using AutoDock. A total of 100 independent docking runs were carried out, and the 15 most populated clusters with the lowest predicted binding energies were selected for further analysis (Fig. 3A). Mapping these conformations onto the three-dimensional structure of MICU1 revealed that the predicted binding pockets were distributed across four distinct regions of the protein (Suppl. Fig. 3). However, because the structural model used for docking corresponds to a MICU1 homodimer, these regions can be grouped into two symmetry-related binding zones. An overlay of the 15 highest-confidence docking poses illustrates this distribution, with regions 3 and 4 representing the symmetric counterparts of regions 1 and 2, respectively (Suppl. Fig. 3). This analysis, therefore, narrows the potential CGP7930 binding interface to two principal regions within the MICU1 structure. One binding pocket is located at the cleft between the C- and N-lobes of MICU1, and the second binding pocket is located between the bridge and the first α-helix of the N-lobe (Fig. 3B).

**Fig. 3.**
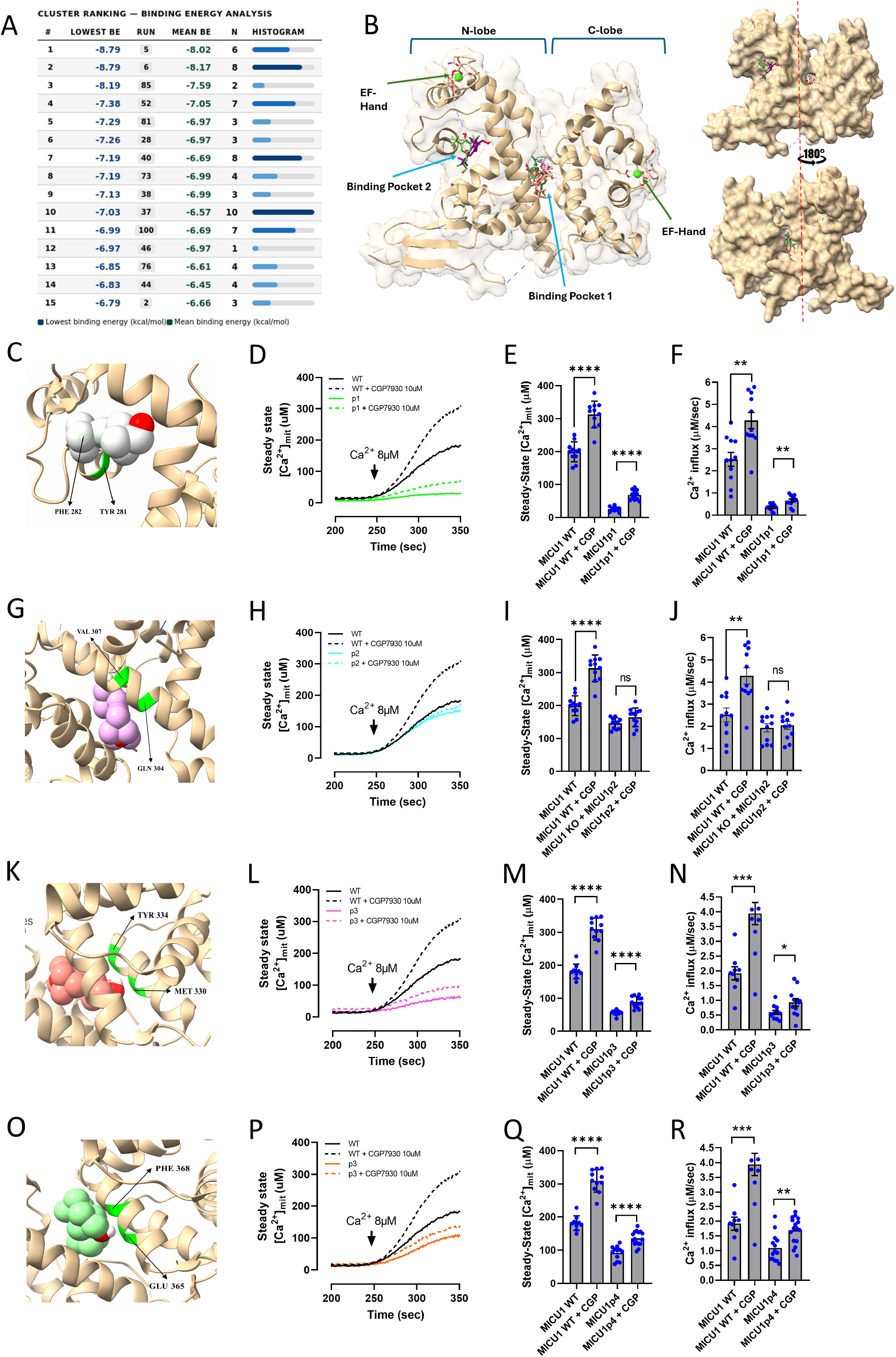
Identification of the CGP7930 interaction site within MICU1 by molecular docking and mutagenesis analysis. (A) Top 15 highest-confidence docking conformations predicted by AutoDock, ranked according to binding energy. The table includes docking run number, mean binding energy, and cluster population frequency for each predicted binding geometry. (B) Structural overlay of the 15 selected docking conformations of CGP7930 bound to MICU1, revealing the principal predicted interaction regions within the protein structure. (C,G,K,O) Three-dimensional representations of the predicted CGP7930-binding regions within MICU1. Residues targeted for mutagenesis are highlighted in green. The corresponding double substitutions generated were Y281A/F282V (P1), Q304G/V307A (P2), M330A/Y334A (P3), and E365A/F368V (P4). (D,H,L,P) Representative traces of [Ca²⁺]ₘ measured in permeabilized HAP cells expressing wild-type or mutant MICU1 constructs together with mitochondria-targeted double-mutant aequorin. Cells were perfused with 8 μM Ca²⁺ in the absence (solid lines) or presence (dotted lines) of CGP7930. Wild-type rescued cells are shown in black, and mutant MICU1-expressing cells are shown in color. (E,I,M,Q) Quantification of steady-state mitochondrial Ca²⁺ levels in permeabilized HAP cells in the absence or presence of 10 μM CGP7930. (F,J,N,R) Quantification of the initial mitochondrial Ca²⁺ uptake rate under the indicated conditions. Data are presented as mean ± s.e.m. from 7-16 independent experiments per condition.

For each docking configuration, the amino acid residues most likely to interact with CGP7930 through hydrogen bonding or hydrophobic/aromatic contacts were annotated. From this analysis, we identified the residues most frequently involved across the 15 highest-confidence docking conformations (Fig. 3A). Eight residues were consistently enriched: Tyr281, Phe282, Gln304, Val307, Met330, Tyr334, Glu365, and Phe368. To experimentally validate these predicted interactions, we designed a targeted mutagenesis strategy. Residues were grouped in pairs based on spatial proximity within the predicted binding regions and substituted with amino acids expected to disrupt the identified interactions. The following pairs of mutations were generated and verified by sequencing: Y281A/F282V (P1) (Fig. 3C), Q304G/V307A (P2) (Fig.3G), M330A/Y334A (P3) (Fig.3K), and E365A/F368V (P4) (Fig.3O). P1/P2 localized in the first binding pocket and P3/P4 localized in the second binding pocket (Fig.3B).

To functionally validate the predicted interaction sites, mitochondrial Ca²⁺ uptake was measured in MICU1 KO HAP1 cells expressing the indicated double substitutions (P1–P4), permeabilized and perfused with 8µM Ca^2+^. The primary objective was to determine which mutations attenuate or abolish the effect of CGP7930 on mitochondrial Ca²⁺ uptake. CGP7930 retained its stimulatory effect in cells expressing the P1 (Y281A/F282V) (Fig. 3C-F), P3 (M330A/Y334A) (Fig. 3K-N), and P4 (E365A/F368V) (Fig. 3O-P), producing fold inductions comparable to those observed in cells expressing wild-type MICU1 and indicating that these residues are not essential for compound activity. In contrast, CGP7930 failed to increase mitochondrial Ca²⁺ uptake above vehicle-treated conditions in the P2 mutant (Q304G/V307A) (Fig. 3G-J), identifying this two-residue region as critical for compound function. Importantly, the Q304G/V307A variant preserved mitochondrial Ca²⁺ uptake capacity comparable to that of wild-type MICU1, indicating that these substitutions do not impair basal MCUc function. By contrast, the P1, P3, and P4 mutants exhibited reduced mitochondrial Ca²⁺ uptake, suggesting partial disruption of MICU1 cooperativity or gating activity. Taken together, these results indicate that MICU1-Gln304 and - Val307 are fully required for the effect of CGP7930 on MCUc activity and suggest that both residues are key determinants for CGP7930 binding to MICU1.

### Analysis of electron density between the residues Gln 304, Val 307, and MICU1

To describe at a fundamental level the nature of the weak interactions between the CGP7930, MICU1-Gln 304, and MICU1-Val 307 in the docking areas, the electron density topology has been analyzed for the most stable docking arrangements that indeed involve both amino acids. To do so, the Non-Covalent Interactions (NCI) index, based on the Reduced Density Gradient (RDG) analysis, was developed by Johnson et al^37^ has been performed.

Figure 4A shows the NCI results for the first arrangement obtained from the molecular docking involving the amino acid GLN304. Firstly, we will focus the discussion in the interaction presented by the OH groups. The OH group belonging to the aromatic ring shows a H-bond with an O atom of the GLN304 amino acid. This bond is represented not only by the corresponding BCP (dotted bond path) but also by the NCI isosurface, which shows a high negative value of sign(λ2)ρ (blue). The other OH group of the isopropyl group forms the same H-bond with the O atom of the TYR334 amino acid. In addition, we found several isosurfaces with negative values close to zero (pale green), which correspond to very weak interactions and are related to dispersion forces between non-polar moieties. In addition, it is worth noting some interactions. That is, some H atoms of the CGP7930 present isosurfaces are associated with H atoms of the MICU1 structure, with high negative values of sign(λ2)ρ. The H···H interactions for nearby hydrogen atoms have been corroborated in bibliography^38^, furthermore the contribution for the stabilization of this kind of interactions has been measured in biological systems with the Molins–Espinosa–Lecomte (EML) approximation^39^, and the results indicate that such H···H interactions contributes significantly to the stabilization^40^, such kind of interaction are presented between the CGP7930 and H atoms of different amino acids (TYR334, GLN304, and TYR369).

**Fig. 4.**
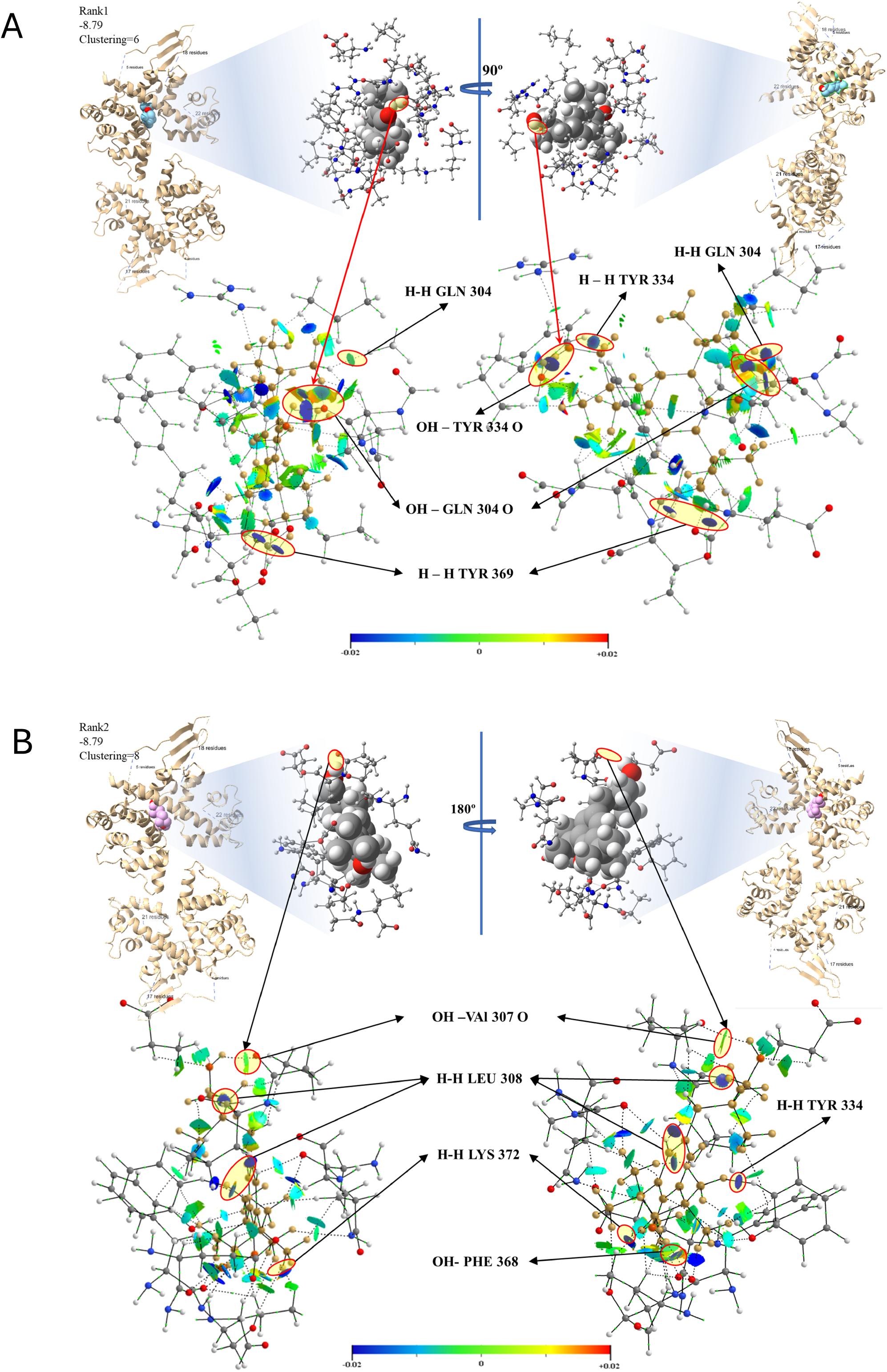
Non-covalent interaction (NCI) analysis of the predicted CGP7930–MICU1 binding modes. To show a clear picture from the trimmed structure used to obtain the wave function, only the interacting moieties have been plotted. Moreover, non-covalent interaction (NCI) isosurfaces were plotted only in the areas between the CGP7930 and MICU1 surrounding the BCPs, avoiding those that appear between the plotted amino acids and would overload the graphical representation. The gradient isosurfaces have been represented using a color mapping scheme that accounts for the values of sing(λ2)ρ. Isosurfaces with blue and pale green, with negative values, are associated with stabilizing interactions, while yellow and red, positive values, correspond to repulsive interactions. Such repulsive interactions are associated with electronic steric crowding and are commonly observed at the centers of aromatic rings and between neighboring functional groups with unshared electron pairs76. Because only the environment between CGP7930 and MICU1 has been plotted to present a clear scheme, no isosurfaces with positive values are present that would appear between stacked amino acids. It should be noted that the NCI index, as determined by the reduced density gradient, is a reliable and consistent method. (A) NCI plot showing the gradient isosurfaces (s = 0.5 a.u.) calculated for the trimmed structure corresponding to the highest-confidence docking geometry identified in the molecular docking analysis. (B) NCI plot showing the gradient isosurfaces (s = 0.5 a.u.) calculated for the trimmed structure corresponding to the second predicted docking geometry. The isosurfaces highlight the non-covalent interaction networks stabilizing the predicted CGP7930–MICU1 complexes.

Regarding the second geometry provided by the molecular docking analysis, a similar scheme is presented. Fig. 4B shows that the OH from the aromatic ring (down) points the H atom towards the PHE368 amino acid, forming an H-bond, as depicted by an isosurface with a high negative value of sign(λ_2_)ρ. On the other hand, the OH group on the aliphatic chain (up) is oriented toward the O atom of the VAL307 amino acid, but in this case, it should be noted that the isosurface is shown in pale green, indicating a less stable interaction. This can be inferred from the distance between the two nuclei. Moreover, in this geometry, in addition to very weak interactions (pale green), isosurfaces with high negative value for sing(λ_2_)ρ between neighboring hydrogen atoms are observed, H···H interactions, can be seen for Leu308, Lys372, and Tyr334 residues.

The computational analyses, including molecular docking and Non-Covalent Interactions (NCI) index based on the peaks that appear in the reduced density gradient (low values of electron density), predicted a favorable binding mode for CGP7930 within a pocket defined by Gln304 and Val307. Consistent with this prediction, substitution of these residues with non-polar amino acids abolished the biological activity of CGP7930. Together, these results support a model in which CGP7930 is stabilized by hydrogen-bond interactions between its hydroxyl groups and residues in the Gln304-Val307 region, thereby promoting its functional effect on MICU1.

### CGP7930 remodels ER Ca^2+^ homeostasis and mitochondrial-ER contact sites

Given the strong cytosolic Ca^2+^ buffering capacity of mitochondria in the presence of CGP7930, it could be expected that a significant dampening of the histamine-induced cytosolic Ca^2+^ transients would occur. However, besides the strong effect of CGP7930 on histamine-induced mitochondrial Ca^2+^ uptake in intact cells, 6-fold increase (Fig. 5A), CGP7930 did not impinge on cytosolic Ca²⁺ transients (Fig. 5B), which agrees with the counter screening evidence (Fig. 1E). This finding raises the question of whether increased mitochondrial buffering capacity of cytosolic Ca^2+^ transients is compensated by the overactivation of alternative cytosolic Ca^2+^ entry pathways. Since IP_3_R is the main contributor to the cytosolic Ca^2+^ peak under histamine stimulation, we measured histamine-induced ER Ca^2+^ release in the absence and presence of CGP7930 in ER-AEQ expressing cells. After ER Ca^2+^ depletion to allow probe reconstitution and subsequent Ca^2+^ refilling, CGP7930 increased histamine-induced ER Ca^2+^ release without altering either the kinetics of ER Ca^2+^ uptake or the resting ER Ca^2+^ levels (Fig. 5C). These results suggest that store-operated Ca^2+^ entry and SERCA activity remain unaffected by CGP7930 and indicate that CGP7930 also promotes directly or indirectly the activity of IP_3_R, a phenomenon previously reported in presence of other MCU activators^41^. To test whether the CGP7930-induced ER Ca^2+^ release requires MCUc activity, we tested the impact of CGP7930 on IP_3_R-driven ER Ca^2+^ release in MCU KD cells (Fig. 5D). In this experimental setting histamine-induced ER Ca^2+^ release was unaffected by the presence of the MCU pharmacological activator, suggesting that the impact of CGP7930 on IP_3_R-driven ER Ca^2+^ release is indirectly mediated by the MCU (Fig. 5D). We propose that increased MCU-driven Ca^2+^ uptake might reduce the magnitude of high Ca^2+^ microdomain generated on IP3R-mouth, thus reducing the well-described negative feedback loop that the IP3R-released Ca^2+^ exerts on its own activity^42^ (Fig. 5F).

**Fig. 5.**
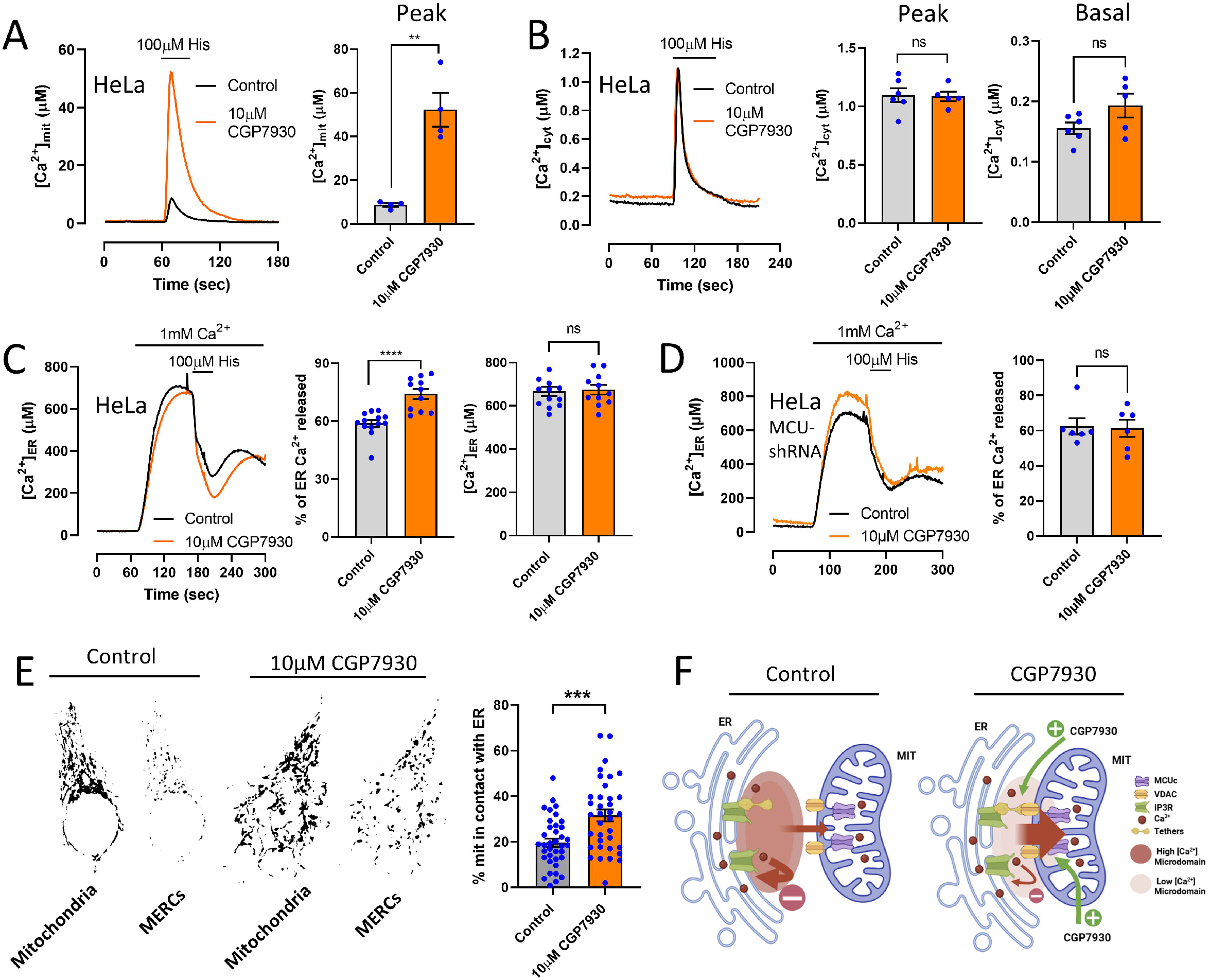
Impact of CGP7930 on cellular calcium remodeling and mitochondria-ER contact sites. (A) Effect of 10µM CGP7930 on histamine-induced [Ca²⁺]ₘ in HeLa cells expressing mitochondria-targeted single-mutant aequorin (mit-119mutAEQ) and reconstituted with coelenterazine w. (B) Effect of 10µM CGP7930 on histamine-induced cytosolic Ca²⁺ concentration [Ca²⁺]c in HeLa cells expressing cytosolic-targeted aequorin (cytAEQ) and reconstituted with coelenterazine w. (C) Effect of 10µM CGP7930 on histamine-induced ER [Ca²⁺]ER release in HeLa cells expressing an ER-targeted double-mutant aequorin (ER-28,119mutAEQ) and reconstituted with coelenterazine w. (D) Effect of 10µM CGP7930 on histamine-induced ER [Ca²⁺]ER release in MCU shRNA HeLa cells. For Ca²⁺ measurements data are presented as mean ± s.e.m. from 4-13 independent experiments per condition. (E) Impact of 10µM CGP7930 on mitochondria-ER contact sites in HeLa cells expressing ER-Mit RspA-split-FAST reconstituted with Lime 4 µM and MitoTracker Deep Red. Data are presented as mean ± s.e.m. from 37-40 cells per condition. (F) Schematic representation of the effects of CGP7930 on ER-mitochondria crosstalk.

Given the low Ca²⁺ affinity of the mitochondrial calcium uniporter^16^, efficient ER to mitochondria Ca²⁺ transfer primarily occurs at mitochondria–ER contact sites^18^ (MERCS). We therefore investigated whether CGP7930 influences MERCS dynamics and promotes close contact formation between the two organelles. To this end, cells expressing the ER–mitochondria Split-FAST reporter and loaded with MitoTracker were treated with 10 μM CGP7930. 1-hour incubation with CGP7930 significantly increased the proportion of mitochondria in contact with the ER (Fig. 5E). These results indicate that, in addition to enhancing MCU activity, CGP7930 also promotes the establishment of ER–mitochondria contacts, thereby potentially facilitating Ca²⁺ transfer between the two organelles (Fig. 5F).

### Pharmacological activation of MCU with CGP7930 boosts excitation-bioenergetics-contraction coupling in the heart

The heart is a highly energy-demanding tissue in which excitation-bioenergetics coupling is tightly regulated. ER to mitochondria Ca²⁺ transfer plays a central role in matching ATP production to the contractile demand of cardiomyocytes^43^. To explore the impact of our new MCU activator on heart function, we performed *ex vivo* perfusion experiments using isolated mouse hearts in a Langendorff system in the absence or presence of 10 μM CGP7930 (Fig. 6A). Left ventricular function was continuously monitored by measuring left ventricular developed pressure (LVDP) and the maximal rate of pressure development (dP/dt). After an initial 5-min stabilization period under control conditions, hearts were perfused with CGP7930 for 15 min. Representative pressure traces (Fig. 6B) show a rapid increase in contractile force and contraction kinetics upon CGP7930 administration. Quantitative analysis of both raw (Fig. 6C and 6D) and normalized (% basal) (Fig. 6E and 6F) data revealed that CGP7930 significantly increased LVDP and dP/dt within 5 min of treatment. Note that these initial experiments were conducted in hearts isolated from C57BL/6 mice. To determine whether these effects were specifically mediated by mitochondrial Ca²⁺ entry through the MCUc, we performed similar experiments using hearts from whole-body MCU knockout mice (CD1 MCU⁻/⁻) and their wild-type controls (CD1) strains. Consistent with our previous observations, CGP7930 induced a significant increase in contractile performance in CD1 control hearts, with elevated LVDP at 5 min and sustained enhancement of dP/dt up to 10 min post-treatment (Fig. 6G-H and 6J-K). Consistently, hearts from MCU-deficient mice failed to respond to CGP7930, showing no significant changes in either LVDP or dP/dt under identical conditions (Fig. 6G,I,J,L). Under none of the experimental conditions, perfusion with CGP7930 altered heart rate (beats per min, BPM), limiting its effect to contractile force (Supplementary Fig. 4). We next evaluated the effects of CGP7930 on mitochondrial Ca^2+^ uptake in cardiac-derived excitable cells (H9C2). Treatment with 10 μM CGP7930 for 1 hour also stimulates vasopressin-induced mitochondrial Ca^2+^ uptake in intact cells (2-fold increase) in a MCU-dependent manner (Fig. 6M-N). These findings demonstrate that the positive inotropic effect of CGP7930 is dependent on a functional MCUc, establishing a direct link between enhanced mitochondrial Ca²⁺ uptake and increased cardiac contractility.

**Fig. 6.**
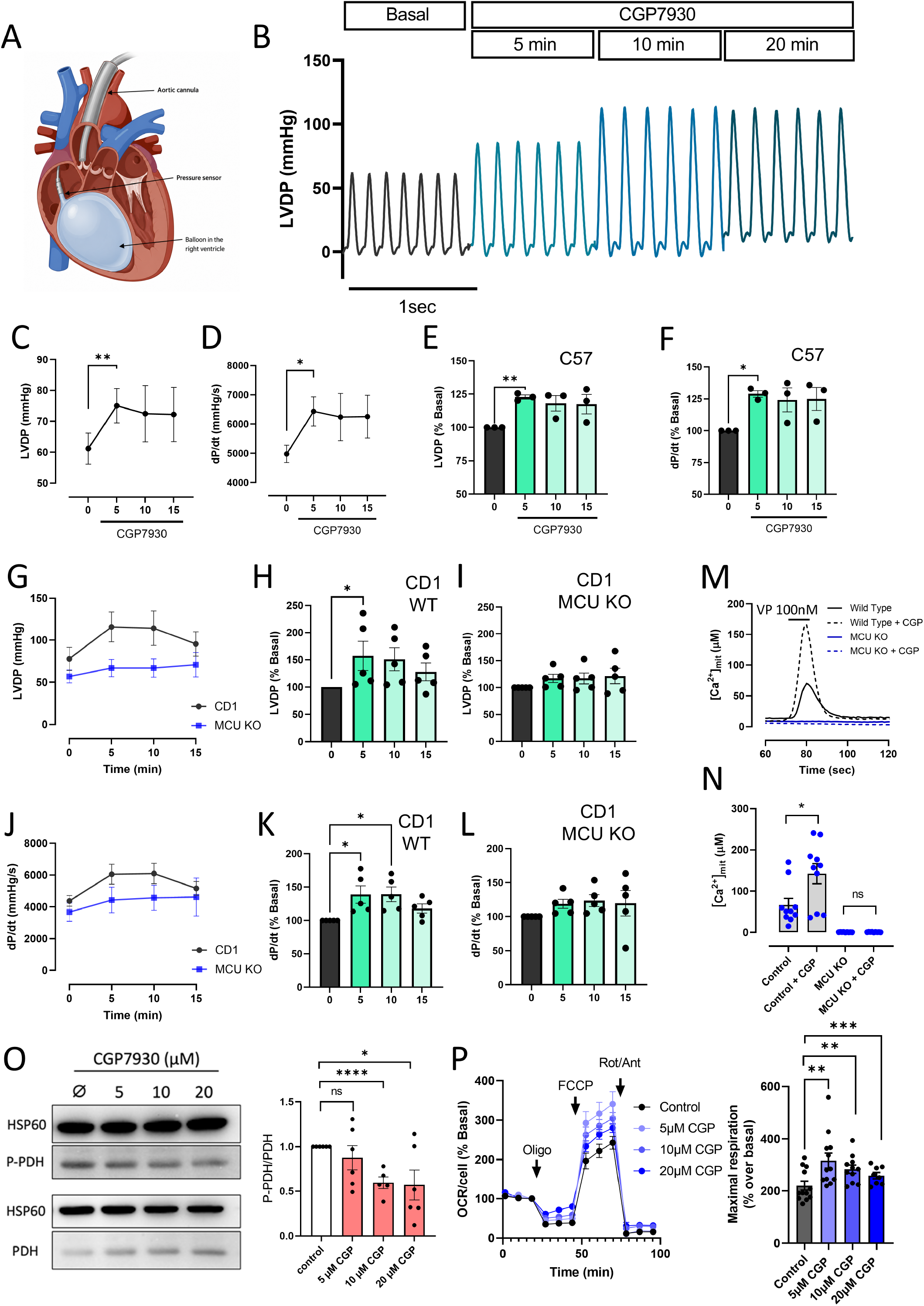
CGP7930 enhances cardiac contractility through MCU-dependent mitochondrial Ca²⁺ uptake. (A) Schematic representation of the Langendorff-perfused heart preparation used for ex vivo functional analyses. The diagram indicates the intraventricular pressure balloon used to measure left ventricular developed pressure (LVDP) and the cannula used for perfusion of Krebs buffer containing CGP7930. (B) Representative pressure recordings from a Langendorff-perfused C57BL/6 mouse heart under basal conditions (black trace; Krebs buffer containing 2 mM Ca²⁺) and following perfusion with Krebs buffer supplemented with 10 μM CGP7930 for 5, 10, and 15 min (dark-to-light green traces, respectively). (C,D) Quantification of raw LVDP (C) and maximal rate of left ventricular pressure development (dP/dt; D) measured in C57BL/6 mouse hearts under basal conditions and after 5, 10, or 15 min of perfusion with 10 μM CGP7930. (E,F) Normalized LVDP (E) and dP/dt (F) values expressed as percentage of basal activity for each heart. (G) Raw LVDP measurements obtained from Langendorff-perfused hearts isolated from wild-type CD1 mice (black) and MCU-knockout mice (blue) before and after perfusion with 10 μM CGP7930 for the indicated times. (H,I) Quantification of normalized LVDP responses in CD1 (H) and MCU-knockout (I) hearts. (J) Raw dP/dt measurements from CD1 (black) and MCU-knockout (blue) hearts under basal conditions and during perfusion with 10 μM CGP7930. (K,L) Quantification of normalized dP/dt responses in CD1 (K) and MCU-knockout (L) hearts. Data are presented as mean ± s.e.m. from 5 mouse hearts per condition. (M,N) Effect of 10µM CGP7930 on vasopressin (VP) -induced [Ca²⁺]ₘ in wild-type and MCU-H9C2 cells. Data are presented as mean ± s.e.m. from 10 independent experiments per condition. (O) Effect of CGP7930 on the phosphorylation status of PDH-Ser293 of ventricular neonatal cardiomyocytes. Data are presented as mean ± s.e.m. from 6 independent experiments. (P) Impact of CGP7930 on uncoupled maximal mitochondrial respiration in ventricular neonatal cardiomyocytes. Data are presented as mean ± s.e.m. from 3 independent experiments and 12 technical replicates per condition.

Mitochondrial matrix Ca^2+^ activates Pyruvate DHase (PDH) via dephosphorylation mediated by allosteric activation of the Ca^2+^-dependent Pyruvate DHase Phosphatase^44^. 10µM CGP7930 incubation of contractile cardiomyocytes significantly reduced PDH phosphorylation by 50% (Fig. 6O), indicative of enhanced PDH activity and increased mitochondrial oxidative metabolism. Consistent with these results, 10µM CGP7930 incubation of contractile cardiomyocytes also enhanced mitochondrial respiration (Fig. 6P). All together, these results are consistent with a model in which CGP7930-driven activation of MCUc enhances the bioenergetic status of cardiomyocytes, thereby supporting increased energetic demand and contractile output in the heart.

## DISCUSSION

Mitochondrial Ca²⁺ uptake through the MCU complex plays a central role in matching cellular energy production to workload and in coordinating intracellular signaling pathways^45^. Despite the fundamental importance of MCU-mediated Ca²⁺ uptake, the number of available pharmacological tools capable of selectively enhancing mitochondrial Ca²⁺ entry remains remarkably limited. In the present study, we performed a high-content screen of 1,280 bioactive compounds and identified CGP7930 as a potent activator of mitochondrial Ca²⁺ uptake (Fig.1). The functional characterization of CGP7930 revealed a robust increase in both the amplitude and rate of mitochondrial Ca²⁺ uptake, both in intact and permeabilized cells (Fig.1F-G; Fig.1H-I). Although originally CGP7930 was described as a positive allosteric modulator of GABA_B_ receptors^46^ and later reported to modulate GABA_A_ receptor activity^47^, it has not previously been linked to mitochondrial Ca²⁺ signaling. In murine models, CGP7930 functions as an anxiolytic and antidepressant, and it has also been demonstrated to reduce self-administration of nicotine, cocaine, and alcohol, although it never progressed into clinical development^48,49^. Our results raise the question of whether the *in vivo* effects of CGP7930 are partially mediated by its impact on MCU activity. Therefore, our screening approach uncovered an unexpected activity for this molecule and establishes CGP7930 as one of the most potent pharmacological activators of MCU-dependent Ca²⁺ uptake reported to date (Fig. 1).

Genetic dissection of the MCU complex demonstrated that the effect of CGP7930 is strictly dependent on the canonical MCU machinery. The complete loss of responsiveness observed in MCU-deficient cells establishes that CGP7930-mediated stimulation of mitochondrial Ca²⁺ uptake requires the pore-forming channel and effectively rules out the involvement of alternative mitochondrial Ca²⁺ uptake pathways. Furthermore, among the regulatory components examined, only MICU1 deficiency abolished the effect of the compound, whereas loss of MCUb, MCUR1, or MICU2 failed to prevent CGP7930-induced mitochondrial Ca²⁺ uptake (Fig. 2). EMRE-deficient cells were not included in our analysis because EMRE is essential for MCU pore-forming subunit assembly, and thus for channel activity^12^. In addition, loss of EMRE disrupts both channel activity and MICU-dependent regulation, making it difficult to distinguish direct effects on MICU1 from secondary consequences of complex disassembly^50^. This selective dependence on MICU1 confirms the gatekeeping machinery of the MCU complex as the main pharmacological target to modulate channel activity^25,31,51^. Interestingly, MCUR1-deficient cells displayed enhanced mitochondrial Ca²⁺ uptake in response to CGP7930. This observation contrasts with the original description of MCUR1 as a positive regulator of MCU activity^15^. However, the precise function of MCUR1 remains controversial, with subsequent studies proposing roles in MCU complex assembly, respiratory chain organization, and mitochondrial bioenergetic homeostasis rather than direct channel regulation^33,52^. One possible explanation for our findings is that loss of MCUR1 induces compensatory adaptations within the MCU regulatory network that increase the sensitivity of the channel to MICU1-mediated activation.

The combined computational and mutagenesis analysis provides mechanistic insight into the mode of action of CGP7930. Although we cannot completely exclude a partial contribution of the P1 (Y281A/F282V), P3 (M330A/Y334A), and P4 (E365A/F368V) regions to compound activity, interpretation of these mutants is complicated by their reduced basal mitochondrial Ca²⁺ uptake, suggesting some degree of MICU1 dysfunction. In contrast, the P2 mutant (Q304G/V307A) preserved basal MCU activity while completely abolishing responsiveness to CGP7930. This selective phenotype provides particularly strong evidence that residues Gln304 and Val307 participate directly in the mechanism through which CGP7930 activates the MCU complex. Consistent with this interpretation, molecular docking identified a preferential binding pocket centered on these residues (Fig.3A and B), and reduced density gradient analysis revealed a network of stabilizing non-covalent interactions involving the hydroxyl groups of CGP7930 (Fig. 4). Notably, previous benchmark studies have shown that NCI analysis is highly robust across computational methods and depends primarily on molecular geometry, supporting the reliability of the predicted interaction network^53^. Interestingly, the proposed MICU1 binding site is very close to those previously associated with MCU inhibitors and activators^25,31^. The convergence of structurally distinct activators and inhibitors on this region suggests that the lobes C and N interface may constitute a privileged regulatory hotspot within MICU1, where small molecules can influence the conformational transitions that govern MCU gating.

The identified binding region offers an attractive mechanistic framework for understanding CGP7930-mediated activation (Fig.3). Structural studies have demonstrated that, under resting Ca²⁺ conditions, MICU1 adopts a conformation that directly blocks the MCU pore through interactions between its conserved polybasic region and the DIME aspartate ring at the channel entrance^54^. Upon Ca²⁺ binding to the EF-hand domains, MICU1 undergoes substantial conformational rearrangements, including compaction of the EF-hand lobes, rotation of the C-terminal lobe toward MICU2, and exposure of hydrophobic surfaces that promote productive interactions with EMRE and channel activation^55,56^. Because the binding pocket identified here is located within the cleft separating the N- and C-terminal lobes of MICU1, we propose that CGP7930 acts as an allosteric activator by stabilizing the Ca²⁺-bound conformation of MICU1. Such stabilization would shift the equilibrium toward the activated state, favoring channel opening even at submaximal cytosolic Ca²⁺ concentrations. This model is also compatible with structural observations showing that MICU1 and MICU2 form a face-to-face heterodimer stabilized by salt bridges and methionine-mediated interactions in the apo state^55,56^. Binding of CGP7930 could facilitate the conformational transition required to overcome this inhibitory arrangement and promote MCU activation.

An unexpected finding of this study was the ability of CGP7930 to increase ER–mitochondria contact sites (Fig. 5E). Because efficient mitochondrial Ca²⁺ uptake relies on the generation of high-Ca²⁺ microdomains at MERCS, this effect could further amplify mitochondrial Ca²⁺ accumulation independently of direct MCU activation. Several non-mutually exclusive mechanisms may explain this observation. First, enhanced mitochondrial Ca²⁺ buffering could reduce the local negative feedback exerted by Ca²⁺ on IP₃ receptors, prolonging Ca²⁺ release events at ER and reinforcing ER–mitochondria communication (Fig.5C and F)^57^. Second, sustained changes in mitochondrial Ca²⁺ signaling may influence the recruitment or stability of tethering complexes that regulate MERCS architecture^58^. More speculatively, MCU activity itself may participate in feedback mechanisms controlling contact-site dynamics. Future studies will be required to determine whether the increase in MERCS is a direct consequence of MICU1 activation or an adaptive response to altered mitochondrial Ca²⁺ handling.

Our findings suggest that pharmacological enhancement of mitochondrial Ca²⁺ uptake has important physiological consequences in the heart. Perfusion of isolated mouse hearts with CGP7930 increased both left ventricular developed pressure and dP/dt, indicating improved contractile performance. Although genetic ablation of MCU produces surprisingly mild cardiac phenotypes under basal conditions^59^, mitochondrial Ca²⁺ uptake is increasingly recognized as an important regulator of energetic adaptation during elevated workload and pathological stress, a relationship consistently supported by our data. A key limitation of this preparation is the inability to directly measure mitochondrial Ca²⁺ in individual cardiomyocytes; however, several lines of indirect evidence allow meaningful conclusions. CGP7930 increases mito-SR contact sites, which would enhance mitochondrial Ca²⁺ uptake during each SR Ca²⁺ release event without requiring a global elevation of cytosolic Ca²⁺, providing a structurally plausible mechanism for the observed energetic boost. Importantly, Nichtova et al.^43^ demonstrated that enforced ER-mitochondria tethering can cause mitochondrial Ca²⁺ overload, but that this is compensated by the formation of mitochondrial nanotunnels that redistribute Ca²⁺ across the network, preventing mPTP opening.

Consistent with this, CGP7930 does not appear to drive Ca²⁺ overload within the approximately 20-minute experimental window, as contractile function is fully preserved with no apparent signs of mPTP opening, suggesting a moderate and physiologically tolerable increase in mitochondrial Ca²⁺ uptake. The fact that CGP7930 did not alter cytosolic Ca²⁺ transients further suggests its inotropic effect is unlikely to result from direct modulation of sarcolemmal Ca²⁺ channels. Nevertheless, given reported activity of CGP7930 at GABA-A and GABA-B receptors^60^, contributions of peripheral GABAergic signaling cannot be completely excluded. While the ex vivo preparation eliminates CNS-mediated autonomic modulation, both receptor subtypes have been described in cardiac tissue and intracardiac neurons, and CGP7930 can directly activate GABA-A receptors independently of endogenous GABA^61^. Whether these mechanisms contribute to the observed effects remains an open question requiring further dedicated investigation.

From a translational perspective, these findings are relevant to one of the most pressing challenges in cardiovascular medicine: limiting injury during ischemia-reperfusion (I/R). Rapid restoration of mitochondrial ATP production upon reperfusion is a central determinant of contractile recovery and cardiomyocyte survival^62^. The ability of CGP7930 to enhance both contractility and ATP production through a physiologically tolerable, self-limiting mechanism, without signs of Ca²⁺ overload or mPTP opening, suggests it could act as a pharmacological preconditioning agent prior to anticipated ischemic events such as cardiac surgery or percutaneous coronary intervention. Future studies in established I/R models will be essential to determine whether these effects translate into meaningful cardioprotection.

In conclusion, our study identifies CGP7930 as a potent activator of mitochondrial Ca²⁺ uptake, reveals MICU1 as a druggable regulatory node within the MCU complex, and provides mechanistic evidence that small molecules can allosterically modulate MCU gating. These findings expand the pharmacological repertoire available to manipulate mitochondrial Ca²⁺ signaling and open new avenues for investigating the therapeutic potential of MCU activation in diseases associated with impaired mitochondrial function and ER–mitochondria communication.

## METHODS

### Cell culture

Wild-type and knocked-out HAP1 cells (purchased from Horizon Discovery) were cultured at 37°C in a humidified atmosphere (5% CO_2_) in IMID cell culture medium (Thermo, #12440-053) supplemented with 10% (v/v) heat-inactivated FBS (Thermo, #A5256701), and 100 µg/ml penicillin/streptomycin (Thermo, #14140-122). HeLa cells (obtained form ATCC) were cultured at 37°C in a humidified atmosphere (5% CO_2_) in DMEM cell culture medium (Thermo, #21885-025) supplemented with 5% (v/v) heat-inactivated FBS, and 100 µg/ml penicillin/streptomycin (Thermo, #15140122). H9C2 cells (obtained form ATCC) were cultured at 37°C in a humidified atmosphere (5% CO_2_) in high glucose DMEM cell culture medium (Thermo, #11995065) supplemented with 5% (v/v) heat-inactivated FBS, and 100 µg/ml penicillin/streptomycin.

### High-content histamine-induced mitochondrial Ca^2+^ uptake screening

The high-throughput screening includes a primary screen and a hit-confirmation screen of mitochondrial Ca^2+^, followed by a secondary screen of cytosolic Ca^2+^. Mitochondrial and cytosolic Ca^2+^ concentrations were measured in intact HeLa cells by using the Ca^2+^-sensitive luminescent probe aequorin targeted to the mitochondria and cytosol, respectively. 150.000 cells/well were seeded in 384-well plates (Corning, #3903) and 15 000 cells/well were seeded in 96-well plates (Corning, #64810), in standard growth medium. After 24 h, HeLa cells were infected with 200 MOI (multiplicity of infection) of the adenoviral vector (Sirion Biotech, Germany) carrying either mitochondria-targeted aequorin for primary and hit-confirmation screening or the cytosolic-targeted aequorin for counter screening. After 48 h, the medium was removed and the cells were incubated with 5 mM native coelenterazine (Biotium, #10110-1) diluted in extracellular buffer (145 mM NaCl, 1 mM MgCl2, 5 mM KCl, 10 mM HEPES, 10 mM glucose, 1 mM CaCl2, pH 7.4) for 2 h at RT in the dark. Then, coelenterazine was aspired and pure compounds of the TOCRISCREN® library (TOCRIS) were added at 10 mM final concentration in DMSO 1 % for 2 h at RT in the dark. Luminescent signal was measured, in basal condition and after stimulation with 100 mM Histamine (Merck, #7125), diluted in aequorin buffer. To calibrate the Ca^2+^ bioluminescence, HeLa cells were semi-permeabilized with 25 mM digitonin (Merck, #11024-24-1) and 10mM CaCl_2_ (Merck, #21115) in the same extracellular buffer. Biolumiscence was detected at the plate readers FLIPR TETRAmax (Molecular Device, USA) and Cytation 3 (Biotek, USA) for 384 well format and 96 well plates format, respectively.

### [Ca^2+^] measurements

HeLa or HAP cells were plated onto 12-mm round glass coverslips in a density of ∼150,000 cells/well and transfected with plasmids encoding cytosolic aequorin (wtAEQ), mitochondria-targeted single-mutant aequorin (mit-119mutAEQ), mitochondria-targeted double-mutant aequorin (mit-28,119mutAEQ), or endoplasmic reticulum (ER)-targeted double-mutant aequorin (ER-28,119mutAEQ). For aequorin reconstitution in cytosolic or mitochondrial compartments, cells were incubated for 1–2 h at room temperature (22 °C) in standard medium containing 145 mM NaCl, 5 mM KCl, 1 mM MgCl₂, 1 mM CaCl₂, 10 mM glucose, and 10 mM HEPES (pH 7.4), supplemented with 2 μM coelenterazine (coelenterazine i or coelenterazine w, as indicated). For ER-targeted aequorin, reconstitution was performed in Ca²⁺-free standard medium supplemented with 10 μM tert-butylhydroquinone (BHQ) to deplete ER Ca²⁺ stores. This depletion step was necessary to prevent premature aequorin consumption during reconstitution. Where indicated, test compounds were included in the reconstitution medium at the specified concentrations. Following reconstitution, coverslips with adherent cells were transferred to the perfusion chamber of a custom-built luminometer for Ca²⁺ measurements.

For experiments in intact cells, coverslips were superfused with standard extracellular medium containing 145 mM NaCl, 5 mM KCl, 1 mM MgCl₂, 1 mM CaCl₂, 10 mM glucose, and 10 mM HEPES (pH 7.4). Intracellular Ca²⁺ responses were evoked by stimulation with 100 μM histamine. For permeabilized cell experiments, cells were first perfused for 1 min with Ca²⁺-free standard medium supplemented with 0.5 mM EGTA. This was followed by a 1-min incubation with intracellular-like medium containing 130 mM KCl, 10 mM NaCl, 1 mM MgCl₂, 0–3 mM potassium phosphate, 0.5 mM EGTA, 1 mM ATP, 20 μM ADP, 5 mM L-malate, 5 mM glutamate, 5 mM succinate, and 20 mM HEPES (pH 7.0), supplemented with 100 μM digitonin to induce plasma membrane permeabilization. Cells were then superfused with digitonin-free intracellular medium for 5–10 min to allow equilibration. Subsequently, mitochondrial Ca²⁺ uptake was assessed by perfusion with intracellular medium containing defined free Ca²⁺ concentrations, prepared using HEDTA/Ca²⁺/Mg²⁺ buffering systems. All experiments were conducted at 37 °C. Luminescence signals were converted to Ca²⁺ concentrations using a previously described calibration algorithm^63,64^.

### *In vivo* assessment of heart function

All experiments were conducted in accordance with the National Institutes of Health Guide for the Care and Use of Laboratory Animals; the protocols were approved by Thomas Jefferson University’s Institutional Animal Care and Use Committee. C57BL/6 mice (12 weeks old) were heparinized by intraperitoneal injection and, after 10 minutes, were quickly euthanized by cervical dislocation. A thoracotomy was performed to expose and cannulate the aorta, followed by perfusion using a Langendorff apparatus. The heart was perfused with Krebs–Henseleit buffer (in g/L: D-glucose 2.0, magnesium sulfate [anhydrous] 0.141, potassium phosphate monobasic 0.16, potassium chloride 0.35, sodium chloride 6.9, sodium bicarbonate 2.1, calcium chloride 0.22), continuously bubbled with 95% O₂ and 5% CO₂ to maintain pH 7.4, at a constant temperature of 37°C. Flow was controlled using a Radnoti minipump and maintained at 12 mL/min. Isolated hearts with an intrinsic heart rate above 320 bpm and developed pressure above 80 mmHg were included in the experiments. To monitor left ventricular (LV) pressure, a latex water-filled balloon connected to a pressure transducer (ADInstruments) was inserted into the LV. The balloon was inflated to achieve an end-diastolic pressure (LVEDP) of 8–10 mmHg. Hearts were perfused for 10–15 minutes for stabilization. After stabilization, hearts were perfused with either Krebs–Henseleit buffer (control) or Krebs–Henseleit buffer supplemented with CGP3790 (TOCRIS #1513) (10 µM) for 20 minutes, while continuously monitoring and recording LV developed pressure (LVDP) using PowerLab. Subsequent analysis was performed using LabChart software. LVDP and dP/dt were calculated at different time points^43,65,66^.

### Mutagenesis

Site-directed mutagenesis of MICU1 was performed using the QuikChange lightning (Agilent, #210519) mutagenesis kit according to the manufacturer’s instructions. The human MICU1 cDNA template plasmid (SC111147) was obtained from Origene. Primers for nucleotide substitutions were designed using the QuikChange Primer Design software provided by the manufacturer. To generate double amino acid substitutions, mutations were introduced sequentially through consecutive rounds of mutagenesis. PCR amplification was carried out under the conditions recommended by the manufacturer. Following amplification, PCR products were transformed into DH5α competent bacteria, plated onto LB-agar plates containing 100 μg ml⁻¹ ampicillin, and incubated for 24 h at 37 °C. Plasmid DNA was subsequently isolated and verified by Sanger sequencing to confirm the presence of the desired mutations and the absence of unintended sequence alterations. Validated MICU1 mutant constructs were co-transfected with mitochondria-targeted double-mutant aequorin at a 1:2 ratio into MICU1-knockout HAP cells. This strategy ensured that mitochondrial Ca²⁺ measurements were selectively performed in cells expressing the rescued mutant MICU1 variants.

### Genetic edition of HAP1 cells

MICU-2 knockout HAP1 cell line were generated using CRISPR/Cas9-mediated genome editing. Single guide RNAs (sgRNAs) targeting the MICU-2 gene were designed using GenCRISPR gRNA Design Tool to target the ACTGAGAAGAGGAAGTCTCG sequence of the exon-2. The following oligos, Fw: 5’caccgACTGAGAAGAGGAAGTCTCG 3’ and Rv: 5’aaacCGAGACTTCCTCTTCTTCAGTc 3’) were synthesized, hybridized and cloned into a pX459 plasmid (Addgene, #62988). Lipofectamine 2000 transfected wild-type HAP1 cells (Horizon) were maintained in IMDM cell culture medium supplemented with 10% FBS and 1% penicillin/streptomycin for 48 hours. Cells were then trypsinized, pelleted, and seeded in selection medium containing 6 µg/mL puromycin for 7 days, with periodic PBS (ThermoFisher, XXX) washes to remove dead cells and facilitate culture expansion. Single-cell clones were isolated using a Cytek Aurora™ cell sorter (Cytek, Fremont, CA, USA; 3-laser configuration), setting one cell per well in 96-well plates. Approximately 50% of the isolated clones were successfully expanded. Once clones reached sufficient confluency, they were trypsinized and further expanded in 60 mm-diameter round cell culture plates (ThermoFisher, #130181). Disruption of the MICU-2 gene expression was confirmed by RT-qPCR. RNA was extracted from seven independent clones using the RNeasy® Mini Kit (Qiagen, #74104), and cDNA was synthesized using the iScript cDNA synthesis kit (Bio-Rad, #1708890), following the manufacturer’s instructions. Quantitative PCR was performed using the Maxima SYBR Green Kit (ThermoFisher, #K0252) on a LightCycler 480 PCR instrument (Roche). The MICU-2 expression was analyzed in each clone using two independent primer pairs targeting the same gene (MICU-2 and MICU-2’) and two independent house-keeping genes (LR18 and ßGUS) to minimize false positives (Supplementary Fig. 2). Primer sequences are listed below.

**Table 1:**
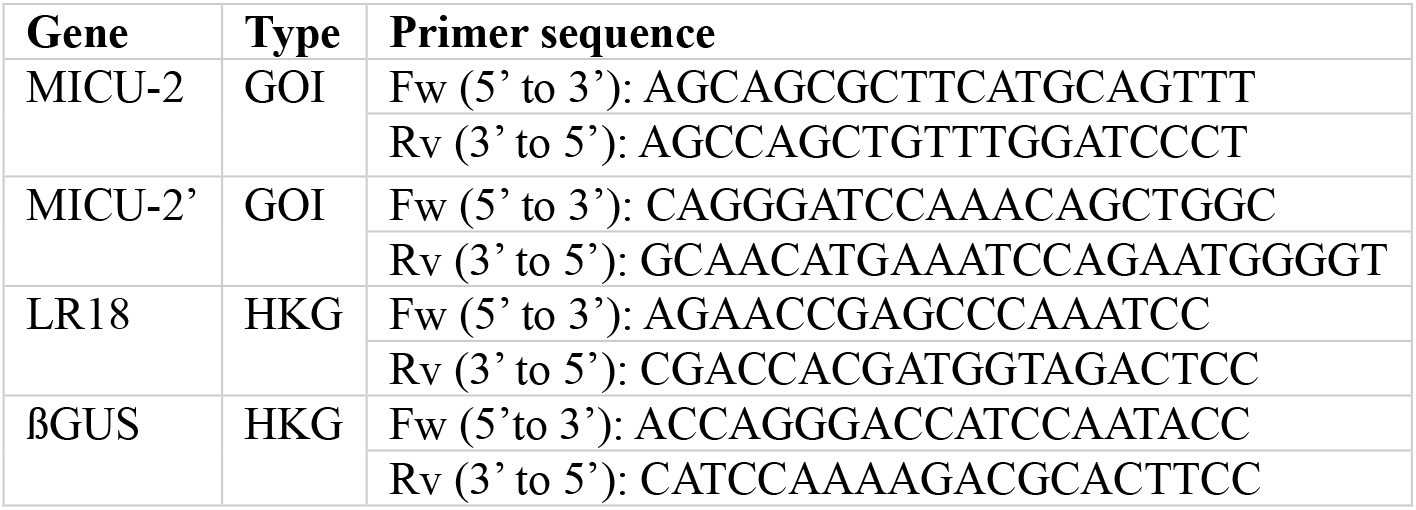
Housekeeping genes (HKG) and genes of interest (GOI) with forward and reverse primer sequences.

### Mitochondrial-ER Contact sites measurements

HeLa cells expressing ER-Mit RspA split-FAST and seeded on 25mm-diameter glass coverslips were loaded with MitoTracker Deep Red (300 nM, Thermo Fisher, #M22426) for 20 min in KRBH (or cell culture medium). After washing twice with KRBH (medium), cells were incubated either with 10µM CGP7930 or DMSO and imaged by confocal microscopy right after reconstitution with the fluorophore Lime (4 µM, Twinkle Factory 480541-250). Confocal imaging was performed on a Leica TCS-II SP5 microscope (Leica), equipped with a 63×/1.4 N.A objective, a tunable WLL laser, and PMT detectors for signal collection. Split-FAST fluorescence was stimulated at 488nm and emission was recorded between 498 and 5300nm. MitoTracker Deep Red fluorescence was stimulated at 644nm and emission was recorded between 655 and 690nm. Images were collected by keeping the same excitation/emission, zoom, and image size parameters. 3D-stacks were collected with *z*-steps of 0.4 µm. Images were background-subtracted and analyzed with specific *ImageJ* (NIH) plugins / background subtracted and preprocessed using a custom-written Python script^67^. The final calculation of the fraction overlap between the MitoTracker Deep Red and the ER-Mit splitFAST signals was performed as described in Garcia-Casas et. al 2024. Briefly, after normalization and binarization of the images, they were merged using pixel multiplication and the fraction overlap of both images was calculated on the MitoTracker image.

### Ventricular neonatal cardiomyocytes isolation

Neonatal ventricular cardiomyocytes were isolated from 1–2-day-old Sprague–Dawley rats (Charles River Laboratories) by enzymatic dissociation. Briefly, ventricles were excised, minced into small fragments, and subjected to serial digestion with collagenase type II (Worthington Biochemical Corporation, #4174) at 37 °C. The resulting cell suspension was collected and enriched for cardiomyocytes by differential adhesion. Cells were initially plated in DMEM supplemented with 5% FBS, 10% horse serum (Thermo, #26050-070), and antibiotics. After 1 h, adherent fibroblasts remained attached to the culture dish, whereas cardiomyocytes remained predominantly in suspension. The non-adherent cell fraction was collected, centrifuged at 700g for 5 min, and resuspended in DMEM containing antibiotics and 2% FBS. Cardiomyocytes were plated onto fibronectin-coated (Thermo #33010018) dishes at a density of 2 × 10⁴ cells cm⁻². Under these culture conditions, cells established intercellular contacts and developed spontaneous contractile activity within 24 h after plating.

### Western-blot

Fibronectin-coated 60-mm diameter Petri dishes were seeded with neonatal ventricular cardiomyocytes at a density of 2 × 10⁴ cells cm⁻² and maintained until they developed spontaneous contractile activity. The day of the experiment, cardiomyocytes were equilibrated in KRBH for 30 min at 37°C. Subsequently, cardiomyocytes were treated with CGP7930 or DMSO (#D12345, ThermoFisher) for 1h. Cell lysis was carried out after 60 min of incubation. Cells were lysed for 15 min on ice in RIPA buffer (Thermo, #89901) supplemented with cOmplete Protease Inhibitor Cocktail (#11697498001, Roche) and phosSTOP phosphatase inhibitor (#4906845001, Roche), for complete protease and phosphatase inhibition. Lysate was centrifuged at 14,000 g for 20 min at 4°C, and the protein content of the supernatant was determined by using Pierce BCA Protein Assay Kit (#23225, ThermoFisher). An amount of 25 µg of total protein was loaded onto SDS-PAGE gels. For immunoblotting, proteins were transferred onto nitrocellulose membrane using a i-blot (Bio-Rad, Spain) and probed with the following antibodies: anti-PDH (#2784S; Cell Signaling), anti-PDH-pSer293 (#ab92696; Abcam), anti-HSP60 (#12165T; Cell Signaling). All antibodies were used at 1/1000 dilution. Horseradish peroxidase–conjugated secondary antibodies (Thermo, #34580) were used 1:10000 in 0,5% BSA (Merck, #A7906-1006) tween-TBS (Biorad, #1706531) for 1 h, followed by chemiluminescence detection (SuperSignal™ West Pic PLUS Chemiluminescent Substrate, Thermo Scientific). Western blots were imaged with a VersaDoc^TM^ 1000 (BioRad) imaging system and quantified with FIJI.

### Oxygen Consumption Rates Measurements

Oxygen consumption in neonatal ventricular cardiomyocytes was measured by using an Seahorse XFe24 metabolic flux analyzer (Agilent). Cardiomyocytes were seeded onto Fibronectin-coated tissue culture plates (#102340-100, Agilent) at a density of 20,000 cells per well. After 2 days, spontaneously contracting cardiomyocytes were washed twice and incubated in DMEM-assay medium (#103575-100, Agilent) supplemented with glucose (#A2494001, Thermo Scientific), Pyruvate (#103578-100, Agilent) and Glutamine (#103579-100, Agilent). Experimental group wells were preincubated with increasing concentrations of CGP7930 (5-30uM) or DMSO in quadruplets. Respiration rates were determined every 6 min. All experiments were performed at 37°C. Respiratory chain inhibitors were added as indicated in the figures at the following concentrations: rotenone (#R8875, Merck), 1 µM; antimycin A (#A8674, Merck), 1 µg/ml; oligomycin (#4110, TOCRIS), 2,5 µg/ml; FCCP (#0453, TOCRIS), 2µM.

### Computational analysis

In order to describe the binding sites between the CGP7930 and the MICU1, we used as starting point the structure from the Protein Data Bank (4NSD), Molecular docking was carried out with AutoDock software^68^, a grid box comprising the entire protein structure was created in a resolution of 0.8 Å. Grid potential maps were computed using AutoGrid 4.2, the docking computation consisted in 100 independent runs. From the 100 docked positions we have considered the most populated clusters with the most stabilizing energy.

In order to gain insight into the nature of the interaction at fundamental level between the compound and the protein in the docking areas, the topology of the electron density has been analysed for the most stable arrangements resulting for the docking analysis. Non-Covalent Interactions (NCI) index, based on the Reduced Density Gradient (RDG) analysis, developed by Johnson et al^37^ has been performed. Such interactions must be considered due to the relevance in chemistry and biology, playing an important role in the interaction between any protein and any drug^69^. The wave function of the resulting geometries found in the molecular docking was generated with the Gaussian 09 package^70^. Since dispersion forces must be considered, the wave function was obtained with the B3LYP-D3 functional^71^, which includes Grimmés dispersion corrections^72^ along with the 6–31 G(d,p) basis set^73^.

AIMAll software package^74^ was used for the analysis of the wave function and the NCI analysis. This approach provides a powerful tool that allows a rich 3D representation of non-covalent interactions with isosurfaces based on peaks that arise in the reduced density gradient at low values of electron density (ρ). The isosurfaces are mapped and colored attending to values of the sign of the second Hessian eigenvalue (λ2). Negative values correspond to stabilising interactions and are depicted in blue and pale green, while positive values correspond to destabilising interactions and are represented in red and yellow. In order to avoid the generation of a huge wave function and carry out affordable DFT calculations, the structure was trimmed by keeping the regions in which the CGP7930 interacts with the surrounding amino acids thorough the so-called Bond Critical Points (BCPs)^75^

## FIGURE LEGENDS

**Suppl. Fig. 1.**
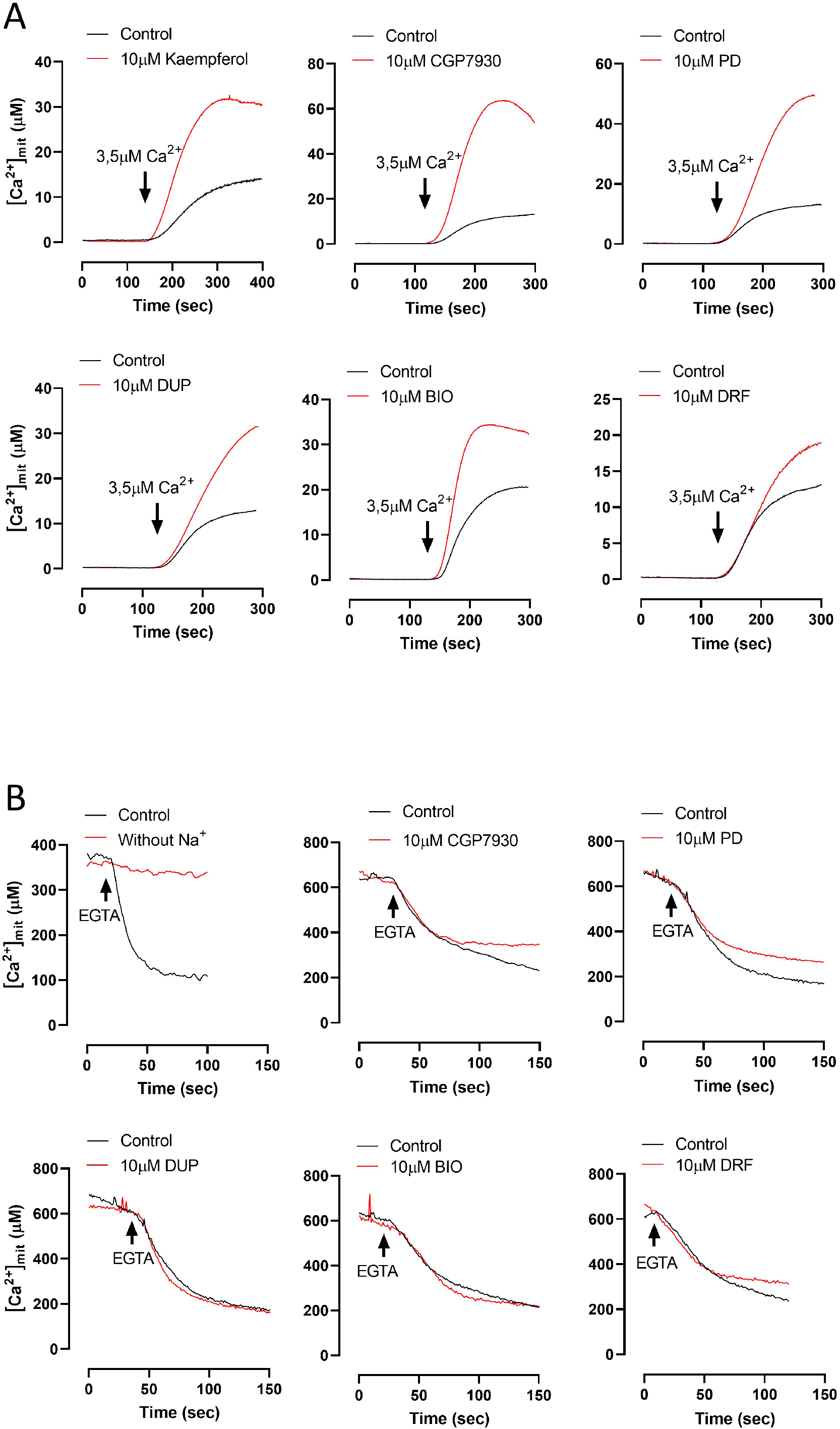
Representative time courses showing the impact of the screening primary hits on mitochondrial Ca2+ uptake in permeabilized cells. Representative time courses of [Ca²⁺]ₘ) measured in permeabilized HeLa cells expressing mitochondria-targeted single-mutant aequorin (mit-119mutAEQ) and reconstituted with coelenterazine w. (A) Mitochondrial Ca²⁺ uptake was induced by perfusion with intracellular-like medium containing 3.5 μM free Ca²⁺. Each panel shows a representative control trace and the corresponding drug-treated condition. Cells were only perfused with the compound during the Ca2+ additions: kaempferol (positive control), CGP7930, PD407824, DuP697, BIO, and (R)-DRF053. (B) Once mitochondrial Ca²⁺ uptake reached a steady-state level, 10 μM EGTA was added to induce Ca²⁺ release. The indicated compounds were present only during the EGTA perfusion phase. Experimental conditions included Na⁺-free medium, CGP7930, PD407824, DuP697, BIO, and (R)-DRF053.

**Suppl. Fig. 2.**
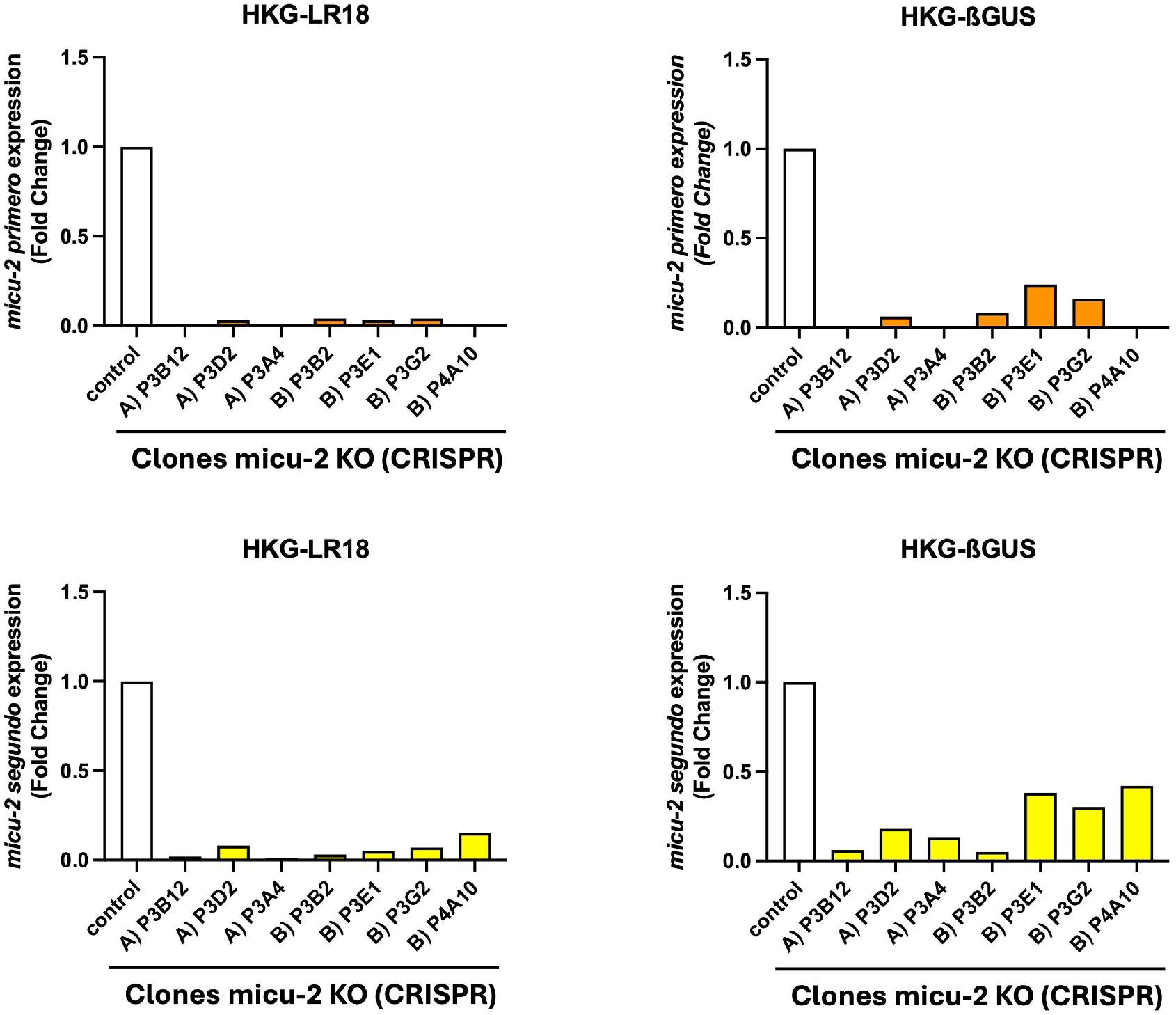
Validation of MICU-2 KO cells. MICU2 mRNA expression in 7 puromycin-selected CRISPR-Cas9-edited HAP1 clones. The results were validated with two different pairs of primers (MICU2-primero and MICU2-segundo, as shown in the upper and lower panels), and two different housekeeping genes (LR18 and ßGUS, as shown in the left and right panels). The clone P3B12 was used in Fig. 2Q-T experiments.

**Suppl. Fig. 3.**
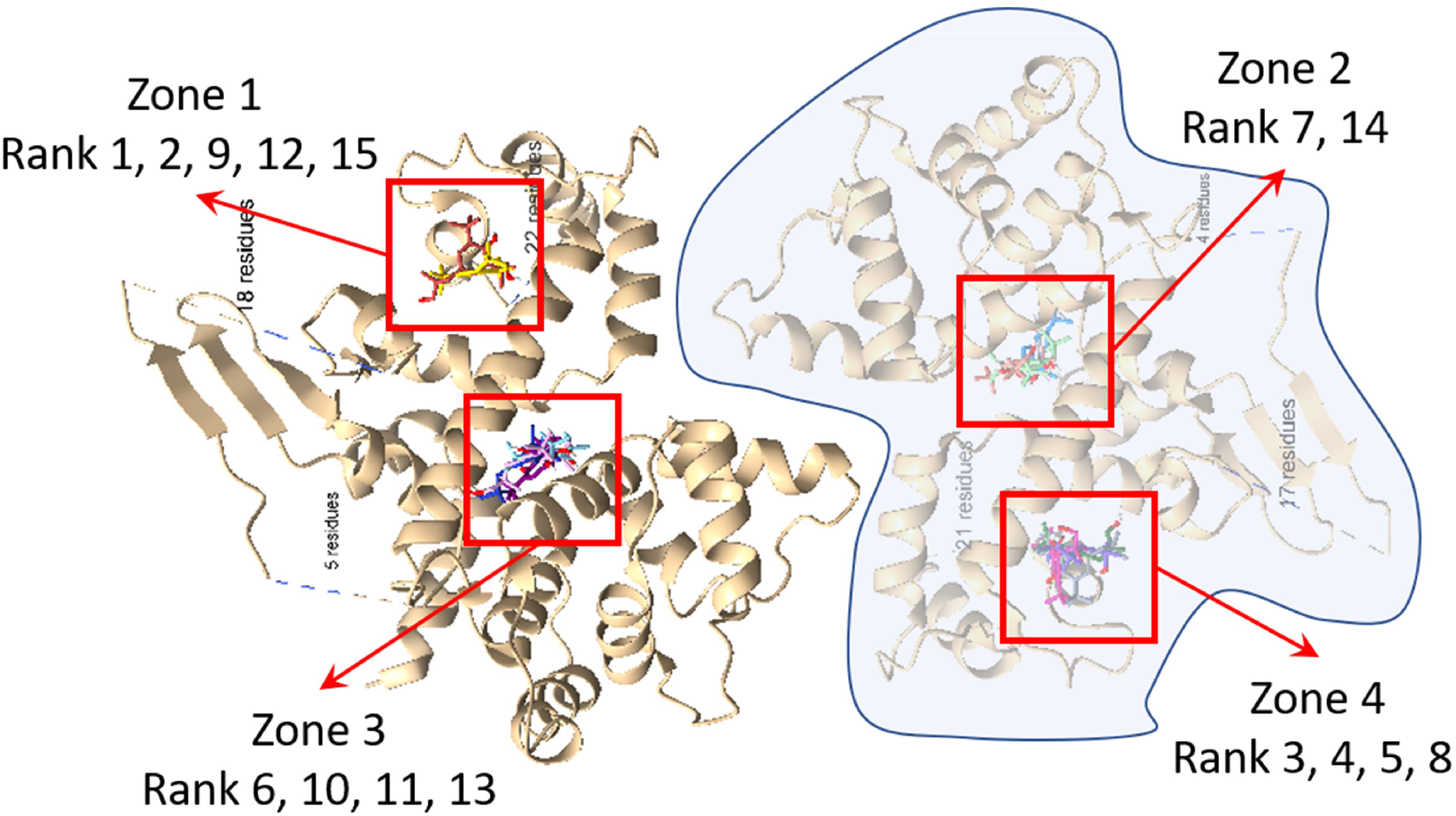
Main binding-sites across the MICU-1 homodimer structure. Mapping the top 15 conformations onto the three-dimensional structure of the MICU1 homodimer (PDB-4NSD) revealed that the predicted binding pockets were distributed across four distinct regions of the protein.

**Suppl. Fig. 4.**
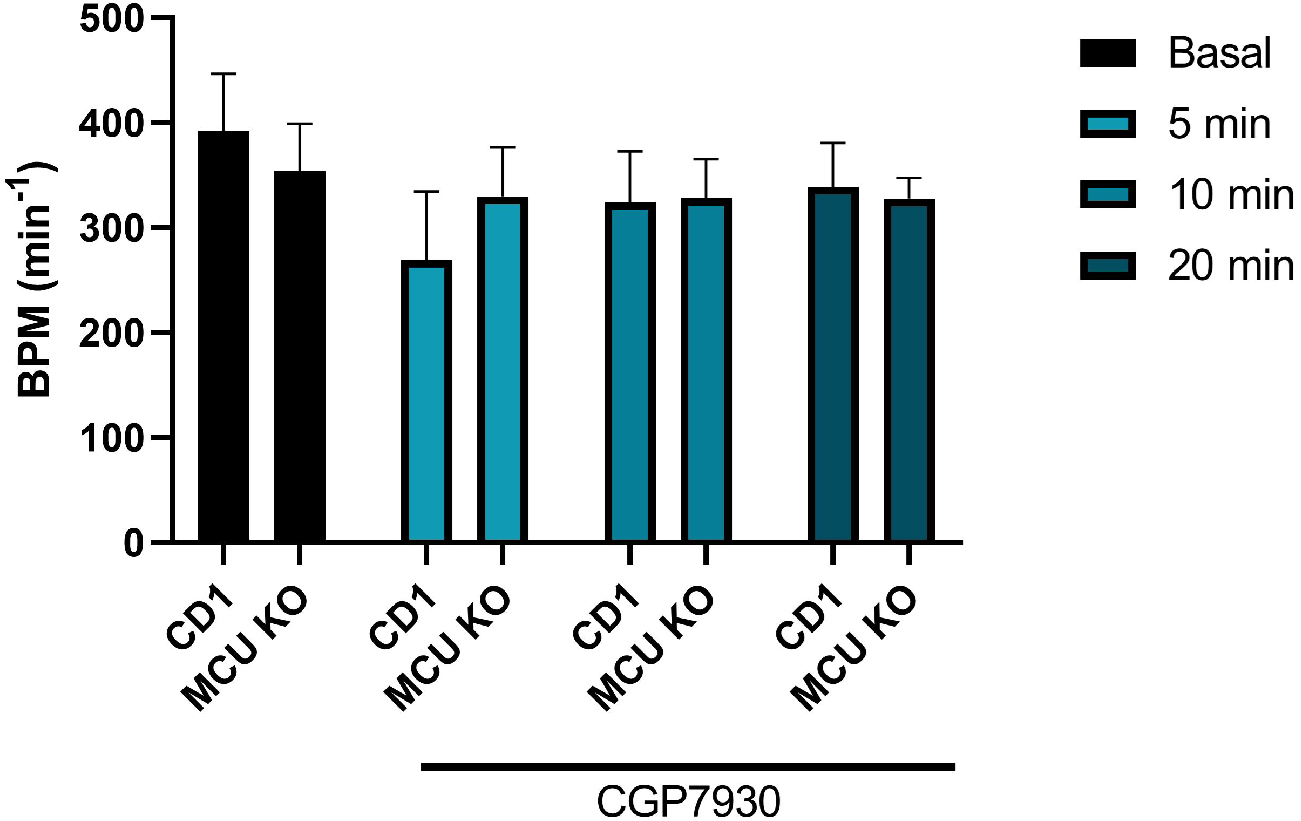
CGP7930 does not alter heart-beating frequency. Bar-chart showing the effect of 10µM CGP3790 on heart beating frequency (Beats Per Minute - BMP) in wild-type and MCU-KO mice hearts at different time points. Data are presented as mean ± s.e.m. from 5 independent experiments per condition.

**Suppl. Table 1.** List of compounds selected after primary reconfirmation and counter-screenings. The table displays the name of the compound, its impact on mitochondrial Ca2+ uptake (AUC), its main defined molecular target and the references supporting the impact on the defined target.

| COMPOUND | AUC (Ca<sup>2+</sup>) | MOLECULAR TARGET | REFERENCES |
| --- | --- | --- | --- |
| (R)-DRF053 dihydrochloride | 1628393 | Potent CK1 inhibitor; also inhibits cyclin-dependent kinases | PMID: 15837193, 16884302, 27210748 |
| PD 407824 | 1695650 | Selective inhibitor of checkpoint kinases Chk1 and Wee1 | PMID: 30234987, 18698753 |
| BIO | 1715731 | ATP-competitive glycogen synthase kinase-3 (GSK-3) inhibitor. Also inhibits cyclin-dependent kinases | PMID: 14700633 |
| DuP 697 | 1899447 | COX-2 inhibitor | PMID: 10542447 |
| CGP 7930 | 2773993 | Positive allosteric modulator of GABAA and GABAB receptors | PMID: 16507713, 16406445, 11641424 |

## Acknowledgments

We thank Niny Rao for facilitating our contact with A. Sánchez-González, who performed all the computational analyses. A. Sánchez-González is especially grateful to Prof. Bruno Victor for his invaluable guidance on docking methodologies, insightful discussions, and continuous support throughout this work. Finally, we thank P. Álvarez-Illera for her excellent technical assistance.

## Author contribution

E.C.E. performed most of the aequorin experiments, as well as the PDH and molecular biology experiments, and analyzed the data with the help of S.R.S and A.M.C; M.F. and S.S.S. performed and analyzed the whole-heart experiments (Nestlé was not involved in mouse study); A.S.G. and A.G. performed the molecular docking and computational analysis; S.F.M. performed the Seahorse experiments; L.G.M and A.A. designed and created the CRISPR/Cas9 plasmids; P.G.C performed and analyzed the ER-MT contact sites microscopy experiments; F.V, B.B, J.N.F and U.M performed the initial compound screening; R.F., M.M., and J.A. revised the manuscript and provided critical feedback; finally, S.F. and J.S.D. designed the experimental approach, performed aequorin and seahorse experiments, analyzed data, and prepared the initial and final version of the manuscript for submission.

